# The *Shewanella oneidensis* Fic enzyme SoFic targets the switch-I region of EF-Tu for AMPylation

**DOI:** 10.64898/2026.07.09.737422

**Authors:** Svenja Runge, Vivian Pogenberg, Alexander Baumgart, Bente Siebels, Hartmut Schlüter, Michael Hecht-Bucher, Aymelt Itzen

**Author notes:** Corresponding Author. (AI.).

## Abstract

Fic enzymes mediate diverse post-translational modifications, including adenosine monophosphate (AMP) transfer and removal, referred to as AMPylation and deAMPylation, respectively. We identified the prokaryotic translation elongation factor Tu (EF-Tu) as an AMPylation target of the Fic enzyme SoFic. SoFic can constitutively reverse EF-Tu modification via deAMPylation whereas AMPylation depends on SoFic homodimerization. The complex crystal structure between SoFic and EF-Tu confirms a conserved target binding mode across evolutionary distant Fic enzymes. AMPylation disrupts EF-Tu’s regulatory switch-I region, causing translational inhibition. SoFic furthermore binds to its promotor DNA, suggesting a dual function as transcriptional and translational regulator in bacterial cells. Together, our structural and biochemical data provide valuable insights into the functional and regulatory diversity of Fic enzymes.

## Introduction

First identified to cause a filamentous phenotype in *Escherichia coli*, filamentation induced by cyclic-AMP enzymes (Fic enzymes) have gained attention due to their presence across all domains of life and because of a remarkably high functional diversity [1–3]. The family of Fic enzymes is defined by the conserved Fic domain, also called Fic/Doc or Fido domain. This domain adopts a characteristic α-helical fold that is shared by Fic enzymes, Doc toxins and AvrB effectors [2].

Many Fic enzymes can use adenosine monophosphate (ATP) as a co-substrate to covalently transfer the adenosine monophosphate (AMP) onto Ser, Thr or Tyr residues of a target protein, resulting in a posttranslational modification (PTM) referred to as AMPylation [4]. Notably, some Fic enzymes can also reverse the modification by deAMPylation using the same catalytic domain [5–7]. Furthermore, a broad range of other PTMs is found among the Fido domain family, including phosphorylation, phosphocholination, UMPylation or rhamnosylation [8–11]. Also the biological contexts, in which these PTMs occur, are diverse: reaching from phage addiction toxin-antitoxin modules or bacterial competition within procaryotes to host cell modulation during bacterial infections and protein homeostasis in eukaryotes [12–17, 10]. Previous studies identified G-proteins as targets for multiple Fic enzymes. G-proteins have central cellular functions in trafficking, cytoskeleton or translation elongation and their modifications thus cause critical cellular consequences [18–20].

In AMPylating Fic enzymes, the Fic domain commonly includes a conserved catalytic H_cat_PFX(D/E)GN(G/K)R sequence motif as a central catalytic element. Moreover, many Fic enzymes contain the auto-inhibitory (S/T)XXXE_inh_(G/N) sequence motif [2, 21]. H_cat_ acts as a general base during catalysis of AMPylation and deAMPylation, thus mutation to alanine renders the Fic enzyme catalytically inactive [22, 23]. E_inh_ is essential for the coordination of a catalytic water molecule during deAMPylation and simultaneously obstructs the active site in a way that prevents AMPylation-competent ATP binding [21, 24, 6, 5, 25]. The location of the auto-inhibitory motif differs between Fic enzymes and is used to distinguish three Fic enzyme classes: in class I Fic enzymes, the inhibitory motif is provided by an interacting peptide, often an antitoxin. In class II and III, the inhibitory motif is located either N-terminal or C-terminal of the catalytic motif, respectively [21]. Class II is the largest subgroup of Fic enzymes and some of their members have been studied in detail, such as the human Fic enzyme FicD. Here, the relief of auto-inhibition is induced by an increased flexibility of E_inh_ upon monomerization of the otherwise dimeric enzyme [26, 27]. There are further examples that suggest an important role of oligomerization for the catalytic activity of Fic enzymes [7, 28]. However, the underlying molecular mechanisms for switching between AMPylation and deAMPylation are highly diverse and remain incompletely understood.

The Fic enzyme SoFic from *Shewanella oneidensis* is largely uncharacterized [29, 30]. The gram-negative bacteria *S. oneidensis* is found in sediments and rarely in clinical isolates. It is of high interest for biotechnological applications due to its capability of extracellular electron transfer [31, 32]. SoFic is a class II Fic enzyme consisting of an N-terminal α-helix, the conserved Fic domain, and a C-terminal winged helix-turn-helix (wHTH) domain [29]. Several similar proteins across many species highlight the conservation of wHTH domain containing Fic enzymes, three of which have been studied in more detail [15, 7, 33]. Previous studies reported auto-AMPylation activity for SoFic and toxicity during heterologous expression in *E. coli* for the deregulated SoFic^E73G^ mutant [21, 24]. Yet, the AMPylation target of SoFic remained unknown.

Here, we identified the elongation factor Tu (EF-Tu) as an AMPylation target of SoFic in *E. coli* and correlated the reported cytotoxicity in bacteria with a downregulation of protein synthesis. We furthermore biochemically characterized SoFic’s catalytic activity and resolved the enzyme-target interaction in a complex crystal structure, contributing to the understanding of the function, regulation and enzyme-target interaction of wHTH domain containing class II Fic enzymes.

## Results

### The domain organization of SoFic indicates multiple functions

In many Fic enzymes, the conserved Fic domain is combined with additional protein domains. These often confer protein-protein interactions and, in some cases, DNA binding [23, 34, 13, 15, 7]. The crystal structures of SoFic wildtype (WT) determined in this study (Fig. 1A) and previously (25, 30) suggest dimerization via its N-terminal α-helix (amino acid (aa) L28-L50) preceding the Fic domain (aa 55-280). C-terminal of the Fic domain the enzyme consists of a wHTH-domain (aa 281-432). The Alphafold database (AFDB) predicts 999 similar proteins with this domain organization, originating from 725 individual species (AFDB50 structures) [35, 36]. The InterPro database annotates 1807 proteins from 1533 species with this domain organization [37, 38].

**Fig. 1:**
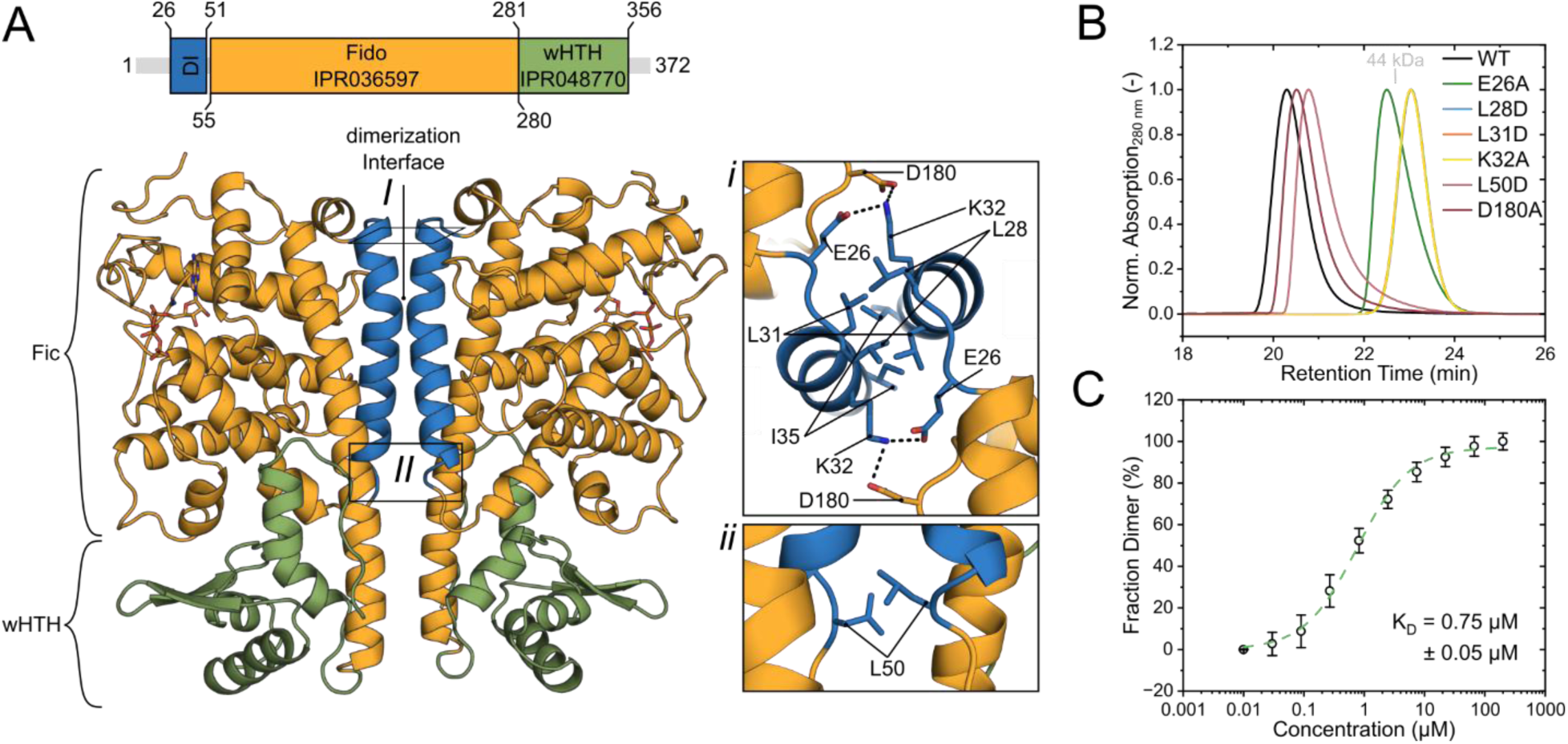
SoFic dimerizes via its N-terminal helix. **A.** Top: Domain organization of SoFic. The Fido domain (green, IPR036597) and protein adenylyltransferase SoFic-like, C-terminal domain (winged-Helix-DNA binding domain, wHTH, yellow, IPR048770) were annotated based on the InterPro entry of SoFic (Q8E9K5)[38, 37]. The dimerizaton interface (DI, blue) was annotated based on the crystal structure of SoFic WT:ATP from this study and previous work [29, 24]. Bottom: Cartoon representation of our SoFic^WT^:ATP crystal structure (ATP is shown as sticks). From the four chains in the asymmetric unit, chains C+D are used for representation. Coloring of the domains according to the schematic above. Inset *I* shows the dimer interface in top view, inset *II* shows a close-up of residue L50 at the N-terminal end of the DI. Potential salt bridges are indicated by dashed lines. **B.** Analytical size exclusion chromatography (aSEC) traces of SoFic^WT^ and dimerization mutants at 50 µM. Each chromatogram was baseline corrected, aligned to an internal standard peak and normalized by the protein peak maximum. The retention time of the 44 kDa reference protein from a gel filtration standard (BioRad) is indicated in grey. **C.** Dimerization affinity of SoFic^WT^ was determined by aSEC using a concentration series of fluorescently labeled SoFic^WT^. The shift of the elution peaks to longer retention times with decreasing concentrations was converted into the fraction of dimeric protein and plotted over concentration. Hyperbolic fitting yielded a K_D_ of 0.75 µM ± 0.05 µM. Error bars represent the standard deviation of two technical replicates for each of two independent biological samples that were labeled with two different fluorescent dyes to rule out dye-related affects on the retention time.

Since additional protein domains can contribute to Fic enzyme function and regulation, we first aimed to identify the functional implications of SoFic’s domain organization. Previous studies reported autoAMPylation of SoFic, indicating AMP-transferase activity of the Fic domain [21, 24]. However, the role of the additional protein domains of SoFic, the N-terminal dimerization helix and the C-terminal wHTH domain, were not addressed experimentally.

We first asked whether the dimerization observed in the crystal structure is transferrable to homooligomerization of SoFic in solution. Indeed, dimerization of wildtype SoFic in solution was observed by analytical size exclusion chromatography (aSEC, Fig. 1B, Table S1): at a concentration of 50 µM, SoFic elutes at an apparent molecular weight (MW) of 90 kDa, indicating dimer formation (calculated MW_SoFic_ = 42 kDa). To estimate the dissociation constant (K_D_) for dimerization, we analyzed a concentration series of 0.01-200 µM SoFic labeled with ATTO-532 or ATTO-488 in aSEC (Fig. S1). Two different fluorophores were used to rule out dye related changes in retention time. With decreasing concentration, the single elution peak of SoFic shifted to longer retention times with a minimal apparent MW of 47 kDa at 0.01 µM. Peak broadening at intermediate concentrations and the absence of two distinct peaks for monomeric and dimeric enzyme indicate a dynamic equilibrium between both species on the aSEC column. We therefore used the apparent MW as a proxy for the fraction of dimeric protein (from 100 % to 0 % dimer at the highest and lowest apparent MW, respectively) and plotted the resulting values over SoFic concentration. Fitting to a hyperbolic function yielded an approximate K_D_ = 0.75 µM ± 0.05 µM for the dimerization of SoFic (Fig. 1C).

The dimerization interface in the crystal structure of SoFic is mainly characterized by hydrophobic interactions between residues L28, L31 and I35 and their equivalents on the second subunit (Fig. 1A*I*). In addition, K32 likely forms a salt bridge with either residue E26 or D180 of the other subunit. E26 is located immediately upstream of the dimerization helix and D180 is part of the Fic domain. On the C-terminal end of the dimerization helix, L50 engages in another hydrophobic interaction with its counterpart (Fig. 1A*II*). As described for structurally similar Fic enzymes before, we implemented single point mutations within the dimerization interface to either disrupt the hydrophobic interactions by inserting a charged amino acid or to remove salt bridge forming residues [7, 33, 15].The dimerization capabilities of these mutants were then compared via aSEC (Fig. 1B, Table S1): at a concentration of 50 µM, the SoFic mutants L28D, L31D and K32A eluted in a symmetric peak at a retention time correlating to 47 kDa and are therefore considered homogeneously monomeric. The elution peak of SoFic^E26A^ is slightly shifted to a shorter retention time (i.e., higher MW = 53 kDa) and peak tailing further indicates a dynamic mixture of monomeric and dimeric enzyme. The mutations L50D and D180A only cause minor changes in retention time compared to dimeric SoFic^WT^, corresponding to 79 kDa and 85 kDa, respectively. Based on these results, K32 likely forms a salt bridge with E26 rather than D180. Loss of this interaction effectively monomerizes the enzyme. Similarly, disruption of the N-terminal hydrophobic patch consisting of L28 and L31 abrogates dimerization of SoFic. In contrast, the C-terminal region of the dimer interface (L50) may be more flexible structurally, allowing for deflection of the inserted charged D50 residues and less repulsion. Overall, the mutational analysis confirms the dimerization interface observed in the crystal structure of SoFic.

We next assessed whether the C-terminal wHTH domain of SoFic (aa 281-432, Fig. 1A) may facilitate DNA binding, which has been observed for wHTH domain containing Fic enzymes before [15, 7]. In analogy to the structurally related Fic enzyme CccR from *Yersinia pseudotuberculosis*, which acts as a transcriptional (auto-)regulator, we hypothesized that SoFic may bind its promotor DNA [15]. On the *S. oneidensis* genome, the gene encoding SoFic (*mloA*) is found on the complement strand and ca. 300 bp separate the ORF from the preceding gene upstream of *mloA.* The 300 bp upstream sequence is likely to contain the promotor for *mloA* transcription and was therefore amplified from the *S. oneidensis* genome for binding studies (hereafter referred to as pDNA, Fig. S2A,B, Table S2).

Qualitative binding studies were performed by aSEC, revealing co-elution of SoFic^WT^ with the pDNA (Fig. S2C-E). In contrast, monomeric SoFic^L31D^ did not show any co-elution, indicating that dimerization may be required for DNA binding by SoFic (Fig. S2C-E). To gain further insights into the sequence specificity of DNA binding by SoFic, the pDNA sequence was divided into three individual 100 bp long fragments (F1-F3, Fig. S2A,B, Table S2). F1 hereby represents the fragment closest to the start codon while F3 is the most distant. The highest co-elution signal was obtained for F1 (Fig. S2F), indicating that SoFic specifically binds to its own promotor region on the *S. oneidensis* genome with a preference for the first 100 bp upstream of the start codon. Since F1 has a substantially lower GC content than F2 or F3 (29 % vs. ca. 40 %), a randomized 100 bp DNA fragment with the same GC content as F1 was included as a control (Table S2). No co-elution was observed for the random DNA fragment (Fig. S2F), confirming the sequence specificity of DNA binding by SoFic.

Taken together, SoFic can dimerize and bind DNA, the latter depending on dimerization. Specific binding to its own promotor DNA suggests a potential role in auto-transcriptional regulation by SoFic, as previously reported for CccR (13). Furthermore, the oligomeric state is a common regulatory mechanism for the catalytic activity of Fic enzymes [26, 28, 27, 7], which will be further analyzed for SoFic below.

### SoFic AMPylates the switch-I region of EF-Tu

To assess the function and regulation of SoFic’s conserved Fic domain, we first attempted to identify its AMPylation target. Substitution of the conserved regulatory glutamate (E_inh_) of class I-III Fic enzymes with glycine (E73G mutation in SoFic) leads to constitutive AMPylation activity [21, 24]. For SoFic and other Fic enzymes, recombinant expression of such mutants results in reduced bacterial growth, suggesting Fic enzyme mediated toxicity caused by AMPylation of bacterial proteins [21]. Hence, we hypothesized that bacterial growth inhibition may be due to SoFic AMPylating a bacterial target protein. To obtain first insights into the spectrum of potentially AMPylated proteins, we overexpressed SoFic wildtype (WT) or the constitutively AMPylation active E73G mutant in *E. coli* as GFP fusion constructs. Western blotting of bacterial lysates with an anti-AMP antibody [39] revealed AMPylated *E. coli* proteins at a molecular weight of 45 kDa (Fig. 2A, top). The fact that AMPylation is observed in pre-induction samples and uninduced cultures may be due to leaky expression, as the fusion protein and the GFP-control are detectable in these samples as well (Fig. 2A, bottom). The disappearance of the AMPylation signal upon induction of the culture may indicate that SoFic^WT^ can also deAMPylate its target protein (Fig. 2A, top, lane 3)[26]. For SoFic^E73G^, also the induced cells exhibit the AMPylation signal at 45 kDa, which is in line with the deAMPylation deficiency of E_inh_ mutants observed for other Fic enzymes [5, 7].

**Fig. 2:**
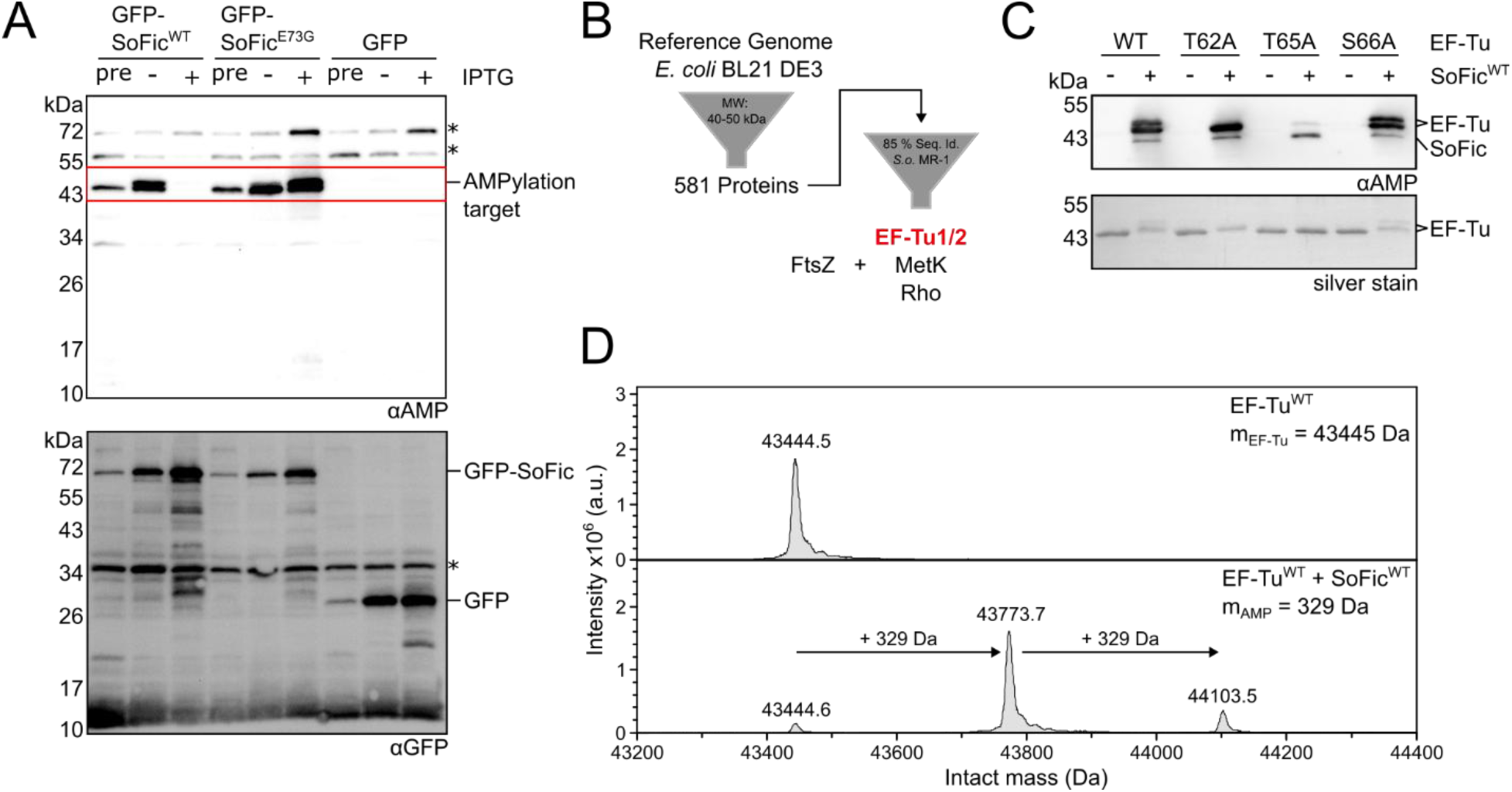
SoFic AMPylates *E. coli* EF-Tu. **A.** Western blot analysis of culture samples after recombinant expression of His-GFP-SoFic constructs in *E. coli* BL21 DE3 overnight. SDS-PAGE samples were normalized by OD_600nm_ of the culture before induction (pre) and after overnight cultivation without (−) or with (+) induction of the expression using 0.5 mM IPTG. Anti-AMP detection (top panel) reveals a SoFic-dependent AMPylation signal at a molecular weight (MW) of 45 kDa. Anti-GFP detection confirms the presence of GFP-fusion proteins and the GFP-control also prior to and without induction, indicating a leaky expression system. Asterisks (*) indicate endogenous background signals in *E. coli*. **B.** Filtering of the reference genome for *E. coli* BL21 DE3 (UniProt UP000002032, Organism 469008) for proteins with a MW of 40-50 kDa and BLASTing (NCBI) of the resulting hits against the genome of *S. oneidensis* MR-I (taxid:211586) suggested three potential AMPylation targets with a sequence identity (Seq. Id.) >85 % between the expression strain and *S. oneidensis*. The candidate FtsZ was added based on previous research [15]. **C.** Western blot analysis of *in vitro* AMPylation of 10 µM *ec*EF-Tu mutants with 0.1 µM SoFic^WT^ and 1.5 mM ATP at 15 °C overnight. 250 ng *ec*EF-Tu were loaded onto SDS-PAGE for Western blotting, anti-AMP detection, and silver staining. **D.** Intact mass spectra of purified unmodified *ec*EF-Tu^WT^ (top panel) and after incubation with 0.1 µM SoFic^WT^ and 1.5 mM ATP at 15 °C overnight (bottom panel). The calculated mass of unmodified EF-Tu (m_EF-Tu_) and the mass shift corresponding to AMPylation (m_AMP_) are indicated.

To identify the AMPylated target protein of SoFic in *E. coli*, we pursued a database research attempt (Fig. 2B, Supplementary Data File 1): Based on the 45 kDa AMPylation signal (Fig. 2A), the reference genome of *E. coli* K12 was filtered for proteins with a molecular weight (MW) of 40-50 kDa, resulting in 581 proteins. We further assumed that SoFic may target an endogenous *S. oneidensis* protein that is conserved in *E. coli* and blasted the *E. coli* hits against the reference genome of *S. oneidensis* MR-1, ranking the results by sequence identity. The *E. coli* proteins with the highest sequence identity to *S. oneidensis* proteins (> 85 %) were MetK, Rho, EF-Tu1, and EF-Tu2. Since EF-Tu1 and EF-Tu2 of *E. coli* only differ by a single C-terminal amino acid, EF-Tu1 was used as a representative in all further experiments. In addition, the cell division protein FtsZ was included as a putative target, because *E. coli* FtsZ is the AMPylation target of the Fic protein CccR, which structurally resembles SoFic and inhibits bacterial growth as well [15]. The four selected target candidates were cloned as His-MBP fusion constructs for co-expression with GFP-SoFic. Again, *E. coli* cultures recombinantly co-expressing SoFic and the putative target proteins were analyzed by Western blotting with an anti-AMP antibody, revealing AMPylation of *E. coli* EF-Tu (*ec*EF-Tu)(Fig. S3).

However, filtering by size and conservation may miss less conserved proteins or those that differ in their molecular weight but do not migrate in SDS-PAGE accordingly. We therefore performed LC-MS/MS analysis of SDS-PAGE gel slices corresponding to the endogenous, AMPylated *E. coli* proteins at 45 kDa in Fig. 2A to assess whether EF-Tu is the predominant AMPylation target of SoFic in *E. coli*. AMPylated peptides with more than one peptide spectra match (PSM) were identified solely for EF-Tu while the few other hits only occurred once, supporting the results from the database approach and co-expression experiment. The AMP moiety on EF-Tu peptides was found at two threonine residues within the switch-I region, T62 and T65, suggesting these residues as modification sites (Supplementary Data Files 2-3, Table S3). Notably, the AMPylation was observed either at T65 (99 % ptmRS localization probability [40]) or at T62 and T65 simultaneously (100 % and 77-92 % ptmRS localization probability, respectively), while AMPylation solely at T62 was not observed.

The same modification sites were also identified in a parallel computational approach: we predicted the complex between one chain of SoFic and one chain of *ec*EF-Tu using AlphaFold2 (AF2)[41]. In the highest ranked interaction model (ipTM = 0.868), the catalytic site was predicted to interact with EF-Tu’s switch-I and revealed the hydroxyl groups of residues T62 and T65, as well as the adjacent S66 in close proximity to H_cat_ (H198) of SoFic (Fig. S4).

To verify the modification sites identified by mass spectrometry and structure prediction, we established *in vitro* AMPylation assays of *ec*EF-Tu using highly purified untagged proteins. Incubation of 10 µM EF-Tu with 0.1 µM SoFic and ATP followed by Western blot analysis with an anti-AMP antibody revealed two distinct AMPylation bands for EF-Tu^WT^ (Fig. 2C top, lanes 1 and 2). The corresponding silver stain furthermore demonstrated a molecular weight shift of EF-Tu due to AMPylation (Fig. 2C bottom, lanes 1 and 2). An additional faint AMPylation signal is observed below the EF-Tu signals, likely corresponding to autoAMPylated SoFic [21, 24], which may not be detectable in the silver stain due to the low concentration. Intact MS analysis of the reaction confirmed the presence of single (mono-) and double (di-) AMPylated EF-Tu (Fig. 2D), likely causing the molecular weight shifts observed in the Western blot analysis (Fig. 2C). Since LC-MS/MS indicated AMPylation of *ec*EF-Tu at T62 and T65, we mutated these residues to alanine (EF-Tu^T62A^ and EF-Tu^T65A^, respectively) and compared the mutants in AMPylation reactions *in vitro* to confirm the modification sites. For EF-Tu^T62A^, only one distinct AMPylation band is observed in the Western blot (Fig. 2C) and homogenously mono-AMPylated EF-Tu is detected by intact MS (Fig. S5). In contrast, the AMPylation signal is substantially reduced for EF-Tu^T65A^ in the Western blot (Fig. 2C) and barely detectable in the intact MS analysis (1.3 % relative intensity for EF-Tu_AMP_, Fig. S5). Based on the AF2 prediction (Fig. S4), also an EF-Tu^S66A^ mutant was analyzed to control for the adjacent S66 as alternative AMPylation site. This mutant was di-AMPylated at levels comparable to EF-Tu^WT^, indicating that S66 is not targeted for AMPylation by SoFic (Fig. 2C, Fig. S5). Together, these results suggest that T65 is the primary AMPylation site within EF-Tu. Moreover, T65 appears crucial for the quantitative AMPylation of EF-Tu by SoFic, since its removal essentially abolishes all considerable AMPylation of EF-Tu. The secondary modification site T62 can only be AMPylated in the presence of T65, while T65 modification does not rely on the presence of T62. However, the data do not allow conclusions on any temporal or causal dependency of the AMPylation at these two residues in EF-Tu^WT^.

We furthermore confirmed that SoFic can also AMPylate *S. oneidensis* EF-Tu (*so*EF-Tu) using co-expression experiments in *E. coli*. Also here, T65 was identified as the main modification site (Fig. S6A). This result is not surprising given the evolutionary conservation of T65 and T62 in EF-Tu. Both residues are found at the C-terminus of the switch-I region of *ec*EF-Tu (aa D51-T65 [42]) and are conserved across the procaryotic EF-Tu superfamily (T62 to 100 %, T65 to 84 %, Fig. S6B). We therefore anticipate that the modification of bacterial EF-Tu by SoFic is species independent due to the high conservation of the AMPylation target.

### SoFic’s AMPylation activity depends on dimerization

Since many Fic enzymes are bifunctional, we analyzed and compared the AMPylation and deAMPylation activities of SoFic using *in vitro* reactions with 0.1 µM SoFic and 10 µM unmodified or AMPylated *ec*EF-Tu^T62A^. The AMPylation reactions were prepared in the presence of 1.5 mM ATP. At specified timepoints, the reactions were stopped by the addition of Laemmli buffer and analyzed by anti-AMP Western blotting. The T62A mutant of EF-Tu was used to avoid double band formation in SDS-Page (Fig. 2C) and simplify densitometric quantification.

AMPylation of EF-Tu by SoFic^WT^ was slow (>240 min, Fig. 3A), while deAMPylation of EF-Tu_AMP_ was completed within 20 min (Fig. 3B). Both catalytic activities of Fic enzymes commonly depend on H_cat_ within the catalytic motif [22, 23, 6, 5], corresponding to position H198 in SoFic. We therefore generated a SoFic^H198A^ mutant, which indeed showed neither AMPylation nor deAMPylation activity towards EF-Tu, confirming the catalytic role of H198 in SoFic (Fig. 3C,D). Furthermore, we repeated the analysis using SoFic^E73G^, the constitutively AMPylation active enzyme. This enzyme revealed a faster onset of detectable EF-Tu AMPylation but yielded lower amounts of AMPylated EF-Tu compared to SoFic^WT^ (Fig. 3E). This result contrasts previous observations of enhanced (auto-)AMPylation activity as well as toxicity of SoFic^E73G^ and the E_inh_ mutants of other Fic enzymes [21, 24]. The lack of any deAMPylation activity of SoFic^E73G^ (Fig. 3F) confirms the catalytic role of E73 during deAMPylation of EF-Tu. Thus, the modification and demodification of EF-Tu by SoFic likely follow the conserved catalytic mechanisms of Fic enzymes [23, 22, 5, 25, 6].

**Fig. 3:**
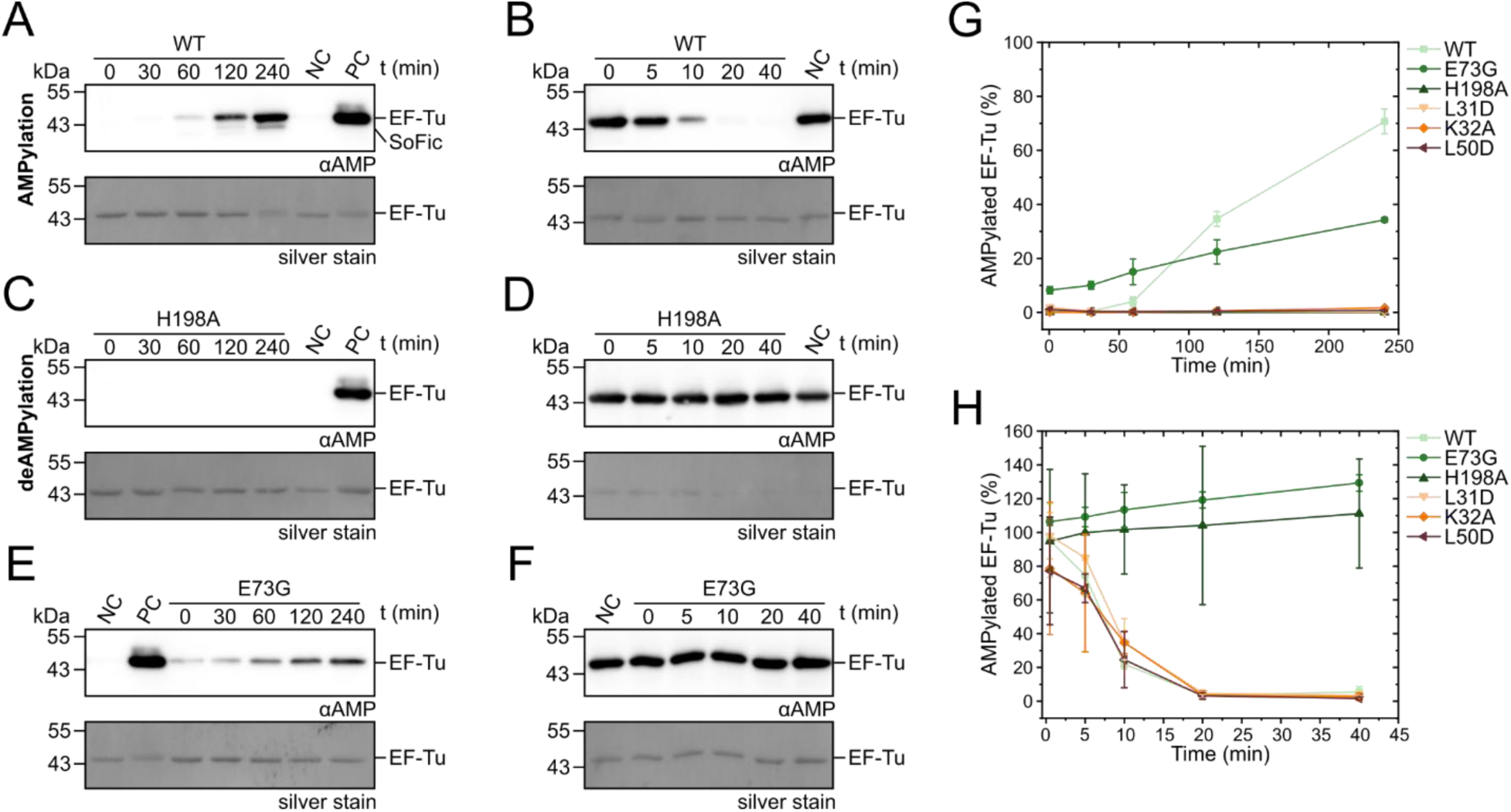
AMPylation of EF-Tu by SoFic depends on dimerization. **A.** Western blot analyses of an *in vitro* AMPylation reaction with 10 µM *ec*EF-Tu^T62A^ and 0.1 µM SoFic^WT^ in the presence of 1.5 mM ATP. At the indicated timepoints, the reactions were stopped by the addition of Laemmli buffer. 250 ng EF-Tu were loaded onto SDS-PAGE for Western blotting, anti-AMP detection and silver staining. NC: negative control without the addition of SoFic (t = 240 min). PC: positive control, 100 % AMPylated *ec*EF-Tu^T62A^_AMP_. **B.** Western blot analyses of an *in vitro* deAMPylation reaction with 10 µM *ec*EF-Tu^T62A^_AMP_ and 0.1 µM SoFic^WT^. At the indicated timepoints, the reactions were stopped by the addition of Laemmli buffer. 250 ng EF-Tu were loaded onto SDS-PAGE for Western blotting, anti-AMP detection and silver staining. NC: negative control without the addition of SoFic (t = 40 min). **C. -F** Western blot analyses of *in vitro* AMPylation and deAMPylation reactions as in **A** and **B**, using SoFic^H198A^ (**C**,**D**) or SoFic^E73G^ (**E**,**F**). **G.** Densitometric quantification of the AMPylation activities of the indicated SoFic mutants based on Western blot analyses (Fig. 3A,C,E and Fig. S7A-C). The error bars represent the standard deviation between two replicates using purified SoFic from two independent biological samples. **H.** Densitometric quantification of the deAMPylation activities of the indicated SoFic mutants based on Western blot analyses (Fig. 3B,D,F and Fig. S7D-F). The error bars represent the standard deviation between two replicates using purified SoFic from two independent biological samples.

Several Fic enzymes are regulated in their AMPylation and deAMPylation activity by their oligomeric state [28, 7, 26, 27, 43]. We therefore analyzed the effect of the dimerization interface mutations L31D, K32A, and L50D (Fig. 1) on the catalytic activities of SoFic. Both monomerizing mutations L31D and K32A depleted the AMPylation activity of SoFic, suggesting that dimerization may be required for this catalytic activity (Fig. 3G, Fig. S7A,B). Intriguingly, SoFic^L50D^ showed the same AMPylation deficiency as the monomeric mutants (Fig. 3G, Fig. S7C), although the mutant is capable of dimerization (Fig. 1B). The deAMPylation activity remained unaffected in all three interface mutants (Fig. 3H, S7D-F). We further compared the AMPylation of EF-Tu for all available interface mutants after incubation overnight (Fig. S7G). After this prolonged incubation time, minor amounts of AMPylated EF-Tu were detected for the SoFic mutants L31D, K32A and L50D as well as E26A and L28D. However, the signals did not reach levels comparable to SoFic^WT^. The dimeric interface mutant SoFic^D180A^ AMPylated EF-Tu comparable to SoFic^WT^. Thus, monomerization of SoFic as well as disturbances at the C-terminal end of the dimerization helix (L50D mutation) result in AMPylation deficiency, potentially indicating an allosteric connection between the dimerization interface and the active site.

This hypothesis is supported by the fact that the dimerization helix transitions into the inhibitory helix of SoFic (aa Q54-E73) directly downstream of L50, separated by a short loop (Fig. S7H). The inhibitory helix features the conserved inhibitory motif (SSEIE_inh_N, aa 69-74) and the inhibitory glutamate E_inh_ (E73 in SoFic), which sterically prevents AMPylation competent binding of ATP in Fic enzymes [21, 24]. We determined the crystal structures of SoFic^L31D^ and SoFic^L50D^ in complex with ATP (Fig. S8, Table S4) to elucidate whether mutations within the dimer interface may affect the position of E73 or the nucleotide conformation in the catalytic center of SoFic in a way that may cause the AMPylation deficiency of the mutants. To provide a reference structure under similar conditions, we determined the crystal structure of SoFic^WT^ independently from previously published structures (Fig. 1A, Fig. S8, Table S4) [29, 24]. As expected, SoFic^WT^:ATP and SoFic^L50D^:ATP form dimers in the crystal. In both cases, the asymmetric unit contains two dimers that are rotated by ca. 180 °. For SoFic^L31D^:ATP, two subunits are observed in the asymmetric unit that do not form any crystal contacts via the N-terminal dimerization helix of SoFic. The comparison of the crystal structures of SoFic^WT^ and the L50D mutant does not reveal substantial changes of the structure or the dimer interaction in the mutant (root mean square deviation (RMSD) = 0.379 Å, Fig. S8A). The overall structure of monomeric SoFic^L31D^ equals the monomeric subunits of the WT structure as well (RMSD = 0.571 Å, Fig. S8B). Mutations in the dimerization interface therefore do not introduce noticeable changes within the overall crystal structure of SoFic.

A more detailed analysis of the catalytic centers revealed that the sidechain orientation of most catalytic residues (aa H198-R209) and the conformation of the bound ATP molecules are similar across all three SoFic:ATP structures (Fig. S8C). The comparison with the crystal structure of SoFic^E73G^:ATP (PDB: 3ZEC (28)) furthermore shows that the ATP in our structures is bound in an AMPylation non-competent conformation (Fig. S8C). The only minor difference between the SoFic:ATP crystal structures is found in the conformation of residue D202, which is involved in the coordination of the Mg^2+^ ion that is essential for catalysis. Exclusively in the crystal structure of SoFic^WT^ the D202 sidechains of one of two dimers point towards the γ-phosphate of ATP (Fig. S8D), as observed in the SoFic^E73G^:ATP crystal structures (PDB: 3ZEC [24], Fig. S8C) or our EF-Tu:SoFic complex (see below, Fig. 5D). This minor conformational change may indicate that a rearrangement of D202 and thus a contribution to Mg^2+^ ion coordination is more likely in SoFic^WT^ than in the L50D or L31D mutants. Nevertheless, the dimer interface mutations L31D and L50D overall do not cause substantial structural changes within the catalytic center of SoFic. During crystallization, SoFic may be forced into a conformation that does not necessarily reflect the distribution of the enzyme’s structural states in solution. This hypothesis is supported by the observation that the mutated D50 residues of SoFic^L50D^ are separated by less than 3 Å despite expected repulsion of the negative charges (Fig. S8E). We therefore conclude that the crystal structures do not offer a conclusive explanation for the AMPylation deficiency of the monomeric and dimeric interface mutants. Nevertheless, our experimental results indicate that dimerization is essential for the AMPylation activity of SoFic. Moreover, the deAMPylation activity of SoFic is independent of mutations in the dimerization interface and more efficient than AMPylation.

### AMPylation alters the conformation of the switch-I region in EF-Tu and interferes with translation

Identifying EF-Tu as AMPylation target prompted us to investigate the effects of AMPylation on its structure and function. For the structural analyses, we preparatively AMPylated GDP-bound *ec*EF-Tu^T62A^ using SoFic^E73G^, followed by the removal of SoFic via size exclusion chromatography (SEC). The T62A mutant of *ec*EF-Tu was used to avoid a heterogeneous mixture of mono- and di-AMPylated protein. We then determined the crystal structure of *ec*EF-Tu^T62A^:GDP AMPylated at residue T65 (EF-Tu^T62A^_AMP_:GDP) at 1.2 Å resolution (Fig. 4A, Table S5) and compared this structure with the available structure of the non-AMPylated protein (PDB 1EFC [44]).

**Fig. 4:**
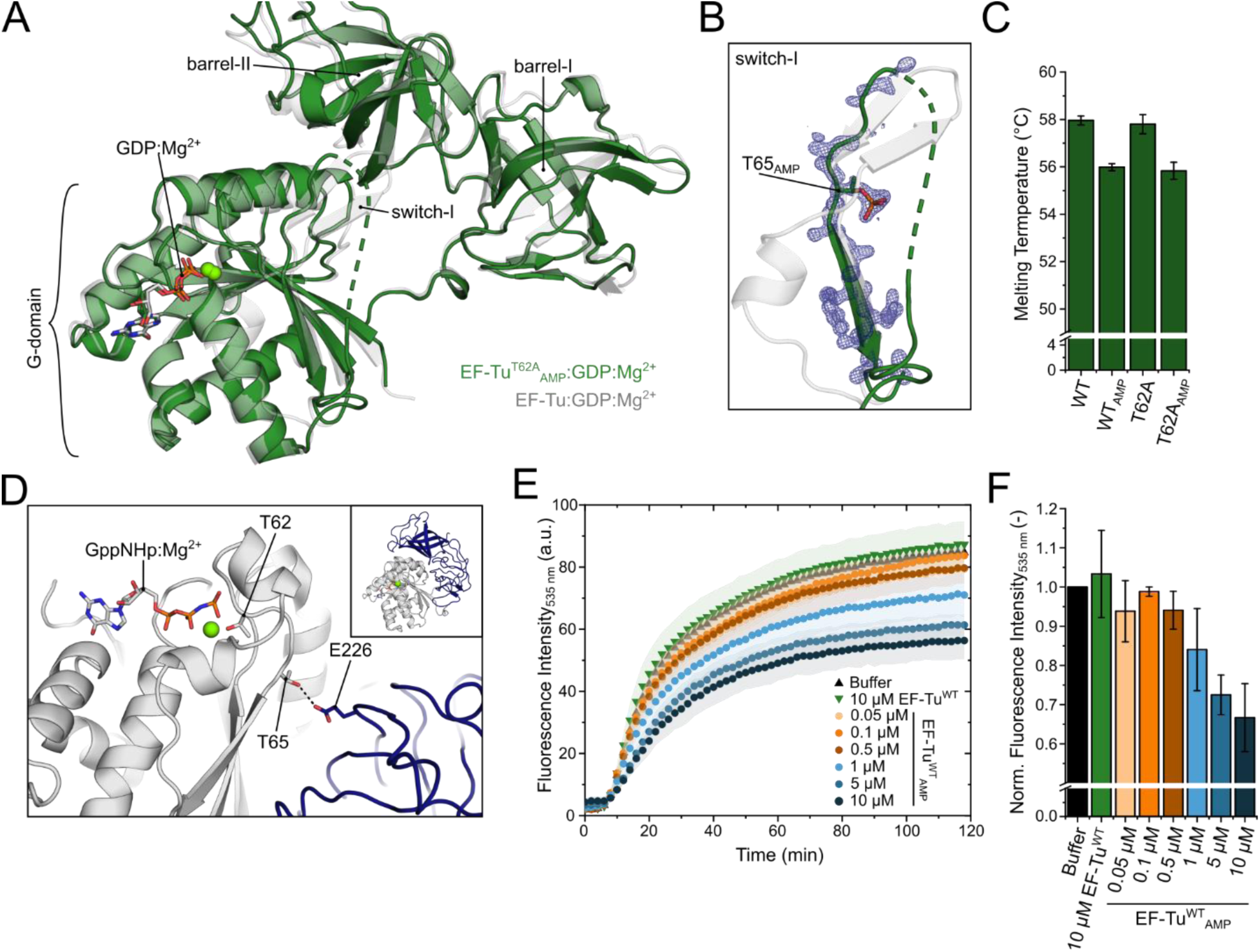
AMPylation by SoFic structurally disrupts the switch-I region of EF-Tu. **A.** Cartoon representation of the crystal structure of *ec*EF-Tu^T62A^:GDP AMPylated at T65 (green) overlaid with *ec*EF-Tu:GDP (transparent grey, PDB 1EFC[44], RMSD = 0.66 Å). Sticks: GDP, green sphere: Mg^2+^. Both structures adopt the open conformation of EF-Tu, where barrel-I is undocked from the G-domain. **B.** Close-up view comparing the switch-I regions of the structures in A. The sidechain of T62 and the α-phosphate of the AMP moiety are shown as sticks. The electron density of AMPylated *ec*EF-Tu^T62A^ is shown as a blue mesh (contour level 2 σ, carve radius 1.6 Å). Coloring as in **A**. **C.** Melting temperatures of unmodified or AMPylated *ec*EF-Tu^WT^ or the T62A mutant were determined by nano differential scanning fluorimetry. Error bars represent the standard deviation of at least three technical replicates for each of two independent biological samples. **D.** Cartoon representation of *T. aquaticus* EF-Tu:GppNHp (PDB 1EFT[45]) showing the interface between the G-domain (grey) and barrel-I (blue), where T65 forms a hydrogen bond with E226 (dashed black line, distance 3.1 Å). The inset at the top right shows the full structure. Barrel-I is docked to the G-domain and the protein adopts a more globular shape compared to the open conformation in A. **E.** Fluorescence of the reporter protein GFP (λ_Ex_=485 nm, λ_Em_=535 nm) over time during coupled *in vitro* transcription and translation reactions in the absence and presence of unmodified or AMPylated *ec*EF-Tu^WT^ at the indicated concentrations. Error bars represent the standard deviation of three technical replicates using purified *ec*EF-Tu^WT^ or *ec*EF-Tu^WT^_AMP_ from two independent biological samples. **F.** Fluorescence intensity values of the last timepoint (t=120 min) in G, normalized to the buffer control. Error bars represent the standard deviation of three technical replicates using purified *ec*EF-Tu^WT^ or *ec*EF-Tu^WT^_AMP_ from two independent biological samples.

EF-Tu consists of an N-terminal G-domain followed by two β-barrel domains (barrel-I and barrel-II). The G-domain contains the nucleotide binding site and the conserved switch-I that is AMPylated by SoFic. In the GDP-bound state, barrel-I is dissociated from the G-domain and EF-Tu adopts an open, elongated conformation. Switch-I (aa D51-T65) forms a short α-helix followed by an antiparallel β-hairpin [44]. The overall conformation of *ec*EF-Tu^T62A^_AMP_:GDP resembles unmodified *ec*EF-Tu:GDP (Fig 4A, RMSD = 0.66 Å) with barrel-I being undocked from the G-domain. The electron density reveals the presence of a phosphate moiety on T65 whereas the adenosine is not resolved, presumably due to high conformational flexibility. Moreover, the β-hairpin that is located immediately upstream of T65 in the unmodified structure (aa E55-N64) as well as the preceding segment including a short α-helix (aa F47-D51) are absent in the electron density of the AMPylated protein. No electron density is observed for most of this region (aa D51-R59), suggesting that it becomes disordered or highly flexible upon modification (Fig. 4B).

To compare the structural results obtained for *ec*EF-Tu to the effect of AMPylation on *so*EF-Tu, we determined equivalent crystal structures of *so*EF-Tu (Table S6). Unmodified *so*EF-Tu:GDP and the T62A mutant both resemble their homolog in *E. coli* (Fig. S9A, RMSD of 0.85 Å and 1.02 Å, respectively), confirming the conservation of the structures of EF-Tu across species. Substitution of T62 with alanine causes no noticeable structural effects. In the crystal structure of AMPylated *so*EF-Tu^T62A^, no electron density is observed for aa F47-S66 including the majority of switch-I and the AMPylation site (Fig. S9B). Likely, the AMP moiety causes conformational heterogeneity of switch-I through steric impairment, thus explaining the lack of defined electron density for this region.

We wondered whether this increased flexibility of switch-I may also indicate a general structural destabilization of EF-Tu. We therefore conducted thermal unfolding analyses of unmodified and AMPylated *ec*EF-Tu and determined the melting temperatures (T_m_) using nano differential scanning fluorimetry (nDSF). The presence of the AMP-group decreased the T_m_ of *ec*EF-Tu^WT^ and its T62A mutant by 2 °C compared to the unmodified proteins (Fig. 4C). Thus, AMPylation at T65 likely interferes with folding of the switch-I in EF-Tu, increasing switch-I conformational dynamics and moderately decreasing the overall structural integrity of EF-Tu.

Since EF-Tu’s activity (e.g., tRNA and ribosome binding), depends on the substantial conformational changes upon GTP binding [42], we also aimed to analyze the structural effect of AMPylation on the GTP-bound conformation of EF-Tu. However, we were unable to obtain diffracting crystals of AMPylated EF-Tu^T62A^ bound to GTP or GppNHp (a non-hydrolyzable GTP-analog), which may be explained by the structural changes caused by GTP binding to EF-Tu: The crystal structure of unmodified EF-Tu from *Thermus aquaticus* bound to GppNHp (Fig. 4D, PDB 1EFT [45]) shows that barrel-I of EF-Tu is docked onto the G-domain, resulting in a compact globular shape. In this conformation, T65 forms a hydrogen bond with E226 (E215 in *ec*EF-Tu) of barrel-I, likely stabilizing the compact conformation. It is plausible that insertion of the sterically demanding AMP moiety at T65 interferes with intramolecular interactions between barrel-I and the G-domain, hence creating a conformationally heterogeneous ensemble of AMPylated EF-Tu molecules that is refractory to diffraction.

During protein synthesis, GTP-bound, domain docked EF-Tu binds amino-acylated tRNA (aa-tRNA) and transports it to the ribosome. Upon recognition of the correct amino acid by the ribosome, EF-Tu hydrolyzes GTP to GDP, transitions into its open conformation and dissociates from the complex, which allows incorporation of the amino acid into the nascent peptide chain. EF-Ts then facilitates nucleotide exchange of EF-Tu and restores active, GTP-bound protein [42]. Since the switch-I region is especially important for EF-Tu’s GDP/GTP-dependent conformational and activity switching, we first analyzed the interactions of EF-Tu with guanin nucleotides and the exchange factor EF-Ts using stopped-flow kinetics as reported previously [46]. However, only moderate changes (factor 2-3 or less) were recorded for the association or dissociation rates of GDP, GppNHp or EF-Ts to AMPylated *ec*EF-Tu compared to the unmodified protein (Fig. S10, Table S7).

We therefore analyzed whether EF-Tu’s function in translation is affected by AMPylation using coupled *in vitro* transcription and translation reactions. We used a commercial cell free expression system that is based on *E. coli* lysates and includes native *ec*EF-Tu. The reactions were prepared using a GFP expression plasmid as DNA template to enable time resolved monitoring of protein expression via fluorescence intensity detection (λ_Ex_=485 nm, λ_Em_=535 nm). The addition of 10 µM unmodified, purified *ec*EF-Tu did not negatively impact the expression of GFP compared to the control reaction (Fig. 4E). Instead, the addition of purified *ec*EF-Tu_AMP_ in concentrations of 0.05 µM to 10 µM reduced the GFP expression up to 30 % in a concentration dependent manner (Fig. 4E,F).

Overall, AMPylation of EF-Tu causes moderate changes in its structural integrity and nucleotide binding properties, while its function in translation is substantially affected. The reduced protein synthesis through AMPylation of *ec*EF-Tu may provide a potential explanation for the previously reported growth defect in *E. coli* during overexpression of SoFic^E73G^ [21].

### Crystal structure of the SoFic:EF-Tu complex

Lastly, we aimed to structurally analyze the enzyme-target interaction between EF-Tu and SoFic. An increased affinity of inactive Fic enzymes to their AMPylated target protein has been reported before [47, 26, 25]. We tested this phenomenon for SoFic and *ec*EF-Tu using aSEC. Co-elution was indeed observed for the catalytically inactive SoFic^H198A^ with AMPylated *ec*EF-Tu^T62A^ (Fig. S11A), but not with unmodified *ec*EF-Tu (Fig. S11B). Based on this observation, we used a 1:1 mixture of SoFic^H198A^ and *ec*EF-Tu^T62A^_AMP_:GDP for co-crystallization and determined the crystal structure of a 2:2 enzyme:target complex at 2.8 Å resolution (Fig. 5A, top, Table S8). Based on the combination of AMPylated *ec*EF-Tu with catalytically inactive SoFic we interpret this structure as a stabilized deAMPylation complex as reported for the eukaryotic Fic enzyme FicD before [25].

**Fig. 5:**
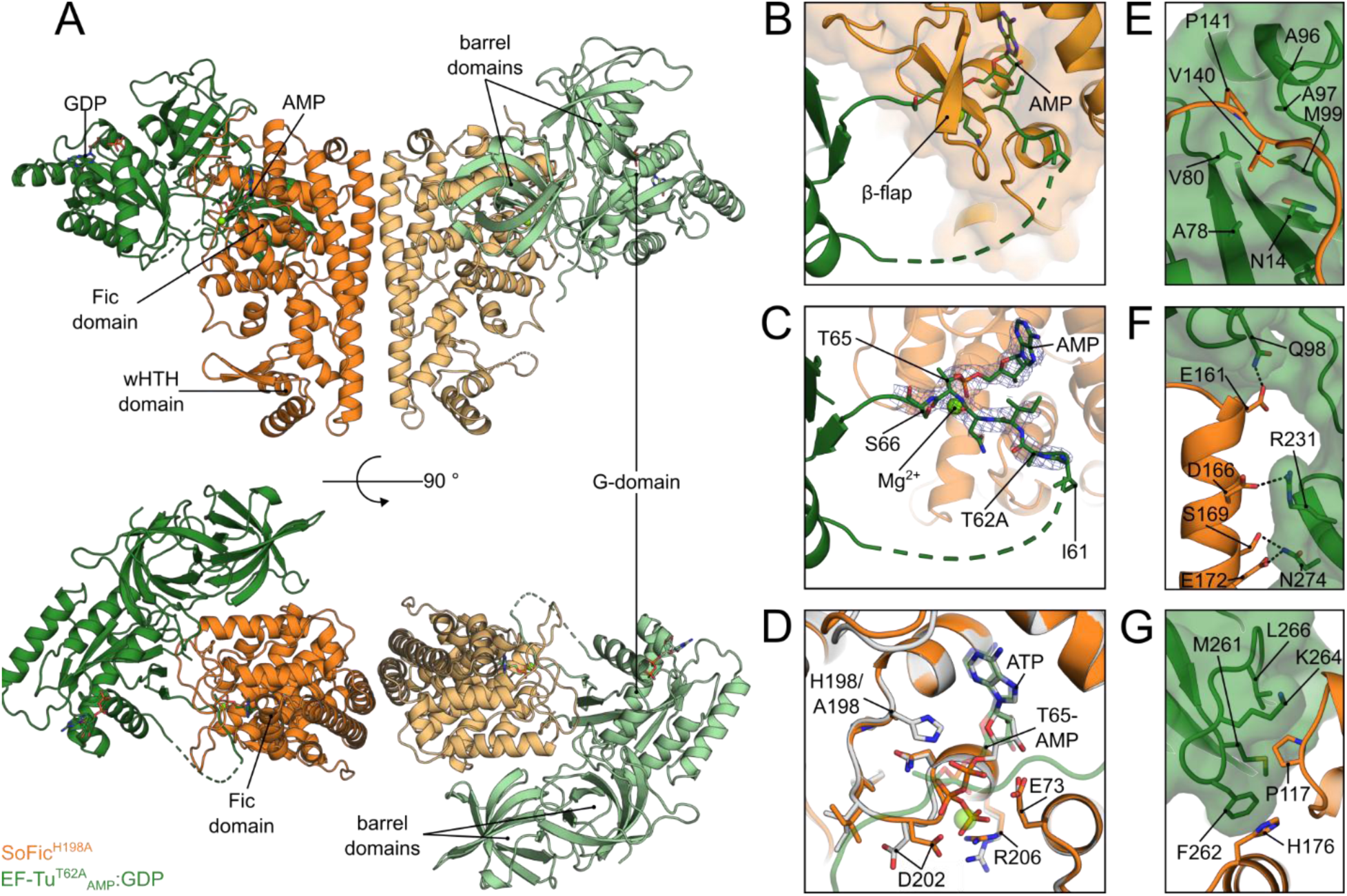
Crystal structure of a SoFic:EF-Tu deAMPylation complex. **A.** Cartoon representation of the complex crystal structure between *ec*EF-Tu^T62A^_AMP_:GDP and SoFic^H198A^ in front view (top panel) and top view (bottom panel). Each subunit of dimeric SoFic^H198A^ (light and dark orange) binds one molecule of EF-Tu^T62A^:GDP AMPylated at T65 (light and dark green), resulting in a 2:2 stoichiometry. **B.** Close up view of the AMPylated switch-I region of *ec*EF-Tu^T62A^, which is rearranged into the active site SoFic^H198A^ and covered by it’s β-flap. The AMP moiety is shown as sticks. Coloring as in A. **C.** Close-up view of the AMPylated region as in B, with the electron density map of aa 61-66 of *ec*EF-Tu as a blue mesh (contour level 2 σ, carve radius 1.6 Å). **D.** Close-up view of the active site of SoFic^H198A^ (orange) binding AMPylated *ec*EF-Tu^T62A^ (transparent green) overlaid with the SoFic^WT^:ATP crystal structure (grey). ATP and AMP are shown as sticks, Mg^2+^ ion of the SoFic:EF-Tu complex is shown as a green sphere. **E.** Hydrophobic interactions between the loop N-terminal of the β -flap of SoFic and the G-domain of *ec*EF-Tu. Coloring as in A, SoFic is shown as cartoon representation, *ec*EF-Tu as cartoon and surface. **F.** Polar interactions between the Fic domain of SoFic and the G-domain (Q98) and barrel-I of *ec*EF-Tu. Dashed black lines indicate hydrogen bonds and salt bridges (distances 2.7-3.4 Å). Coloring as in A, SoFic is shown as cartoon representation, *ec*EF-Tu as cartoon and surface. **G.** Hydrophobic interactions between the Fic domain of SoFic and barrel-I of *ec*EF-Tu. Coloring as in A, SoFic is shown as cartoon representation, EF-Tu as cartoon and surface.

In complex with SoFic, *ec*EF-Tu^T62A^_AMP_ adopts the open GDP-conformation and the GTPase and barrel domains wrap around the Fic domain of SoFic (Fig. 5A, bottom). The modified region of EF-Tu’s G-domain is rearranged into the active site of SoFic and covered by the conserved β-flap of Fic-enzymes (Fig. 5B). Further backbone interactions between the SoFic β-flap and EF-Tu (aa I61-S66) result in β-sheet-like binding of the AMPylated region (Fig. S12A,B). As observed for AMPylated EF-Tu alone, no electron density was obtained for a substantial part of the switch-I region preceding the AMPylation site (aa A43-G60, Fig. 5B,C), suggesting structural flexibility of this region. In contrast to the crystal structure of AMPylated *ec*EF-Tu (Fig. 4A,B), the AMP moiety is fully resolved in the complex crystal structure, likely due to conformational stabilization within the active center of SoFic (Fig. 5C). The position of the adenosine in the active center is comparable to ATP in our SoFic:ATP crystal structure (Fig. 5D). Furthermore, a Mg^2+^ ion is coordinated by D202 of SoFic and the α-phosphate of the AMP-moiety in the deAMPylation complex (Fig. 5C,D).

The interaction between EF-Tu and SoFic is characterized mainly by hydrophobic contacts: in addition to the β-sheet-like interaction, binding in the active site is stabilized by hydrophobic interactions between L145 of SoFic (L145_SF_), Y155_SF_, L244_SF_, L245_SF_, and the adenine base of the AMP-moiety while the ribose forms polar contacts with E73_SF_, E195_SF_, R209_SF_, and Y240_SF_ (Fig. S12C). Moreover, I63 of EF-Tu (I63_EF_) is confined between I154_SF_ and Y241_SF_. Outside of the catalytic center, V140_SF_ and P141_SF_ point into a hydrophobic cleft between A78_EF_, V80_EF_, A96_EF_ and A97_EF_ of the G-domain (Fig. 5E). Further interactions are found between SoFic’s α-helix spanning residues E161_SF_-E172_SF_ and barrel-I of EF-Tu: here, a network of polar interactions stabilizes the enzyme-target complex. This includes hydrogen bonding between Q98_EF_, and E161_SF_, R231_EF_ and D166_SF_, as well as N274_EF_ with both S169_SF_ and E172_SF_ (Fig. 5F). At the C-terminus of this helix, H176_SF_ and F262_EF_ engage in aromatic-aromatic interaction (π–π stacking). Adjacent, P117_SF_ points into a hydrophobic patch between M261_EF_, L266_EF_ and the aliphatic face of K264_EF_ of barrel-I (Fig. 5G). To verify the interfaces observed in the crystal structure, the effects of alanine mutations on the interaction between SoFic^E73G^ and EF-Tu^WT^ were analyzed by *in vitro* AMPylation assays (Fig. S12D). The removal of single hydrophobic residues of SoFic at all of the interfaces described above, namely P117_SF_, P141_SF_, and H176_SF_ (Fig. 5E-G), efficiently reduced the AMPylation of EF-Tu, suggesting that these residues substantially contribute to the enzyme:target interaction. A similar effect was observed upon mutation of I154_SF_ (Fig. S12A), which may be explained either by the weakened interaction with I63_EF_ or by alterations in the β-flap of SoFic, thus disturbing the β-sheet-like stabilization of EF-Tu’s switch-I. Removal of D166_SF_ barely affected the AMPylation of EF-Tu by SoFic, which may suggest that the polar interactions are less important for the interaction or that loss of a single residue can be compensated by the surrounding network of polar interactions. Residue Q250_SF_ was included as a control residue outside of the interaction interface and, as expected, did not affect the AMPylation activity of SoFic^E73G^. Note that we only analyzed the AMPylation activity of SoFic point mutants to validate the interface since mutations of most residues of EF-Tu that are involved in the interaction with SoFic did not yield soluble protein upon recombinant expression in *E. coli*. Nevertheless, the mutational analysis validates that the crystal structure likely represents the interaction of SoFic with EF-Tu in solution. The complex therefore provides valuable insights into the target binding mode during deAMPylation catalysis of SoFic.

## Discussion

In this study we identified the translation elongation factor EF-Tu as a novel AMPylation target of the previously uncharacterized *S. oneidensis* Fic enzyme SoFic. The bifunctional enzyme was shown to be an efficient deAMPylase, while its AMPylation activity depends on dimerization. Specific binding to its own promotor DNA adds additional (auto-)regulatory potential to the enzyme’s function. Our structural and biochemical analyses of AMPylated EF-Tu show that the modification disrupts the conformation of the modified switch-I region, ultimately causing a reduction in protein synthesis. We furthermore resolved a crystal structure of a complex between SoFic and EF-Tu, confirming and extending the knowledge on Fic enzyme-target interactions.

Our data show that SoFic is a bifunctional Fic enzyme, capable of both AMPylating and deAMPylating EF-Tu (Fig. 3A,B). Both catalytic activities of SoFic require H198 (H_cat_, Fig. 3C,D), which is known to act as a general base in Fic enzyme mediated nucleotide transfer [23, 22] The deAMPylation activity of SoFic moreover requires E73 (E_inh_, Fig. 3E,F), which coordinates a catalytic water molecule for the nucleophilic attack on the AMP moiety [5, 25, 6]. The inhibitory glutamate of class I-III Fic enzymes can furthermore prevent binding of ATP in an AMPylation competent manner. Removal of this residue often leads to enhanced AMPylation activity [21, 24, 26, 7, 5, 6]. Intriguingly, the E73G mutant of SoFic did not show the expected enhanced AMPylation activity compared to SoFic^WT^ (Fig. 3E,G). However, it did produce higher levels of AMPylated EF-Tu at early timepoints (i.e. 60 min incubation), which were analyzed in previous studies of SoFic’s autoAMPylation activity [21, 24]. Considering the lack of deAMPylation activity in SoFic^E73G^ and the inhibition of protein synthesis by AMPylated EF-Tu (Fig. 4E,F), the toxicity of this mutant during expression in *E. coli* cells may be caused by the accumulation of AMPylated EF-Tu [21], while SoFic^WT^ appears to deAMPylate EF-Tu after induction of expression (Fig. 2A). How the switch between AMPylation and deAMPylation by SoFic^WT^ is regulated within bacterial cells, however, is not conclusively revealed by our data. We can furthermore not exclude that AMPylation of other proteins in *E. coli* contributes to the toxicity of SoFic^E73G^, since few other AMPylated proteins were detected by LC-MS/MS at low abundance (Table S3).

Some studies have linked the relief of autoinhibition by E_inh_ with monomerization of the respective Fic enzyme [28, 7, 26, 27]. An underlying allosteric mechanism has been described for FicD and is based on an increasing conformational flexibility upon monomerization, causing loss of a hydrogen bond network between the dimer interface and the active center, which allows for the displacement of E_inh_ (E234 in FicD)[26]. In SoFic, the direct transition of the dimerization helix into the inhibitory helix (Fig. S7H) suggests an allosteric connection as well. However, in contrast to FicD, the AMPylation activity of SoFic was not enhanced by monomerizing mutations in the dimer interface. In turn, enforcing monomerization onto SoFic abolished its AMPylation activity without affecting deAMPylation (Fig. 3G,H, Fig. S7). We attribute the contrasting effect of monomerization on SoFic and FicD to the different dimerization interfaces: while SoFic dimerizes via its additional N-terminal α-helix, FicD dimerizes via the Fic domain, rationalizing a differential allosteric signaling.

Since SoFic’s deAMPylation activity was not affected by mutations within the dimerization interface, we propose that E73 of SoFic can adopt the AMPylation inhibiting and deAMPylation active conformation in all oligomeric states, hence SoFic is a canonical deAMPylase. In contrast, AMPylation required an intact dimerization interface. Notably, the *in vitro* reactions in this study were performed at a concentration of 0.1 µM SoFic, corresponding to ca. 10 % dimeric enzyme (0.01 µM). Nevertheless, this small fraction of dimeric enzyme appears sufficient to enable AMPylation, in contrast to the fully monomeric L32D and K32A mutants. We therefore conclude that an allosteric signaling between the dimerization interface and the active center may be required to relieve the active site obstruction by E_inh_ and enable AMPylation. The results obtained for SoFic^L50D^, which is AMPylation deficient despite its ability to dimerize, furthermore support this hypothesis (Fig. 3G, Fig. S7C). The residue is located at the hinge between dimerization helix and inhibitory helix (Fig. S7H). Hypothetically, the hydrophobic attraction between both L50 sidechains upon dimerization may contribute to the rearrangement of E_inh_ and the relief of autoinhibition in the active center of SoFic. This interaction is lacking in SoFic^L50D^ or the monomeric interface mutants L32D or K32A, and may thus inhibit AMPylation.

Comparable to our observations of AMPylation deficiency in monomeric mutants of SoFic (Fig. 3, Fig. S7), the AMPylation dependent toxicity of a SoFic homolog in *E. coli* (*ec*SoFic) was fully depleted upon monomerization. Notably, *ec*SoFic also features a leucine residue (L47) at the position corresponding to L50 in SoFic [33]. In other structurally related wHTH containing Fic enzymes, no corresponding hydrophobic residues in this position were reported, and the AMPylation activity is not affected or enhanced upon mutations in the dimerization interface [15, 7]. Together, these results highlight the diversity of regulatory mechanisms among Fic enzymes, also within structurally related groups, and the need to study individual enzymes to elucidate their respective regulatory mechanisms.

The C-terminal wHTH domain of the aforementioned group of Fic enzymes was previously linked to DNA binding in a transcription factor like manner [15, 7]. In line with CccR from *Y. pseudotuberculosis*, dimeric but not monomeric SoFic specifically binds to its own promotor DNA within the first 100 bp upstream of the start codon (Fig. S2). In its dimeric state, SoFic may therefore act as a transcriptional regulator of its own gene in *S. oneidensis* cells. Although that hypothesis seems plausible in analogy to CccR, the analysis of transcriptional regulation by SoFic was not within the scope of this study [15].

The Fic domain of SoFic catalyzes the AMPylation and deAMPylation of EF-Tu, an essential bacterial G-domain containing protein that is involved in translation elongation during protein biosynthesis. This observation aligns well with previous findings of GTP-binding proteins as central target molecules of bacterial Fic enzymes. Several pathogens secrete Fic domain containing effector proteins into the host cell to hijack its cellular processes, often by the modification of small GTPases: the *Vibrio parahaemolyticus* effector VopS and the *Histophilus somni* effector IbpA both AMPylate switch-I of Rho GTPases to induce the actin cytoskeleton of the host cell to collapse [17, 16]. AMPylation of eukaryotic elongation factors, also consisting of a G-domain has been reported as well, although a physiological relevance of such modifications remains elusive [48, 49].

In bacterial cells, EF-Tu is one of the most abundant proteins and vital for the translation of proteins by transporting aa-tRNA to the ribosome [50, 51, 42]. The large number and high diversity of reported posttranslational modifications of EF-Tu showcase the important role of PTMs to dynamically regulate protein biosynthesis according to changing environmental conditions [52, 53, 20]. To our knowledge, this is the first time that AMPylation of EF-Tu by a Fic enzyme is reported.

The most abundant and best studied modification of EF-Tu is phosphorylation, which also occurs under physiologic conditions [54, 51, 53]. An obvious analogy to our results is the phosphorylation of EF-Tu by the toxin-antitoxin module doc/phd (death on curing/prevent host death) that originates from the prophage P1[14]. Doc belongs to the same Fic/Doc (Fido) superfamily as SoFic’s Fic domain and consists of an incomplete Fido domain fold that is complemented and regulated by Phd [13]. Unlike most other Fido domain enzymes, Doc acts as a kinase and phosphorylates EF-Tu at the conserved T382 (*E. coli*) of barrel-II [11, 55]. Phosphorylation at T382 traps EF-Tu in its open conformation resembling the GDP-bound state independent of the nucleotide bound, which precludes ternary complex formation between EF-Tu:GTP and tRNA and ultimately inhibits translation [11, 20].

Another related example is the phosphorylation of EF-Tu during cellular quiescence of *Bacillus subtilis*. Here, the kinase YabT phosphorylates switch-I at the strictly conserved residue T63 (equivalent to T62 in *E. coli* EF-Tu) [56]. T62 is the secondary AMPylation site of SoFic and close to the main modification site T65. In *B. subtilis* EF-Tu the phosphorylation at switch-I was shown to reduce the GTP hydrolysis rate. Initially, this deficiency was proposed to stabilize the interaction with the ribosome (dissociation of EF-Tu is triggered by GTP hydrolysis) and therefore stalling translation [56, 42]. However, alongside the analysis of EF-Tu phosphorylated at T382 by Doc, analyses of a phosphomimetic EF-Tu^T62E^ mutant indicated that the phosphorylation of switch-I similarly interferes with the conformational change of EF-Tu upon GTP binding, thus precluding aa-tRNA binding and resulting in translation inhibition [20]. Moreover, the previously observed co-precipitation of *B. subtilis* EF-Tu phosphorylated at T62 was linked to a reduced solubility instead [20, 56].

In contrast to the proposed stabilizing effect of T62 or T382 phosphorylation on the characteristic GDP-state β-hairpin conformation of switch-I [20, 57], our results indicate that AMPylation disrupts this structural feature of EF-Tu (Fig. 4A-C). However, AMPylation at T65 is likely to interfere with the docking of barrel-II to the G-domain (Fig. 4D). The effect of AMPylation on the secondary site T62 would furthermore cause severe clashes with the nucleotide and Mg^2+^, if the AMPylated switch-I could adopt the helical conformation that is characteristic for the GTP-bound state (Fig. 4D)[45, 57]. Since the affinity of EF-Tu_AMP_ to GTP is not decreased but rather slightly increased (Fig. S10F), we conclude that AMPylation does not preclude GTP binding but may instead have a similar effect on the conformational equilibrium of EF-Tu than phosphorylation at T62 or T382. This hypothesis is supported by the observation that AMPylated EF-Tu, like phosphorylated EF-Tu, dominantly reduced protein synthesis in coupled *in vitro* transcription and translation reactions despite the presence of native, unmodified EF-Tu in the cellular extract (Fig. 4E,F)[56]. Based on these observations, we assume that EF-Tu_AMP_ still interacts with components of the translation machinery. However, the exact molecular basis for the reduced protein synthesis was not conclusively determined.

The physiological relevance of EF-Tu AMPylation by SoFic will be interesting to investigate in the future. Since endogenous *so*EF-Tu can be AMPylated by SoFic (Fig. S6), the enzyme may be an endogenous tool of *S. oneidensis* to quickly adapt growth rates to changing environmental conditions, as reported for phosphorylation of EF-Tu in *B. subtilis* during sporulation [56]. Another mechanism has been shown for the structurally related Fic enzyme CccR from *Y. pseudotuberculosis*: Here, AMPylation of the cell division protein FtsZ inhibits cell division of competing bacterial species [15]. Notably, also EF-Tu has been identified as an interaction partner of CccR in a pulldown. Since this interaction was not analyzed and we currently do not have information about a potential secretion of SoFic by *S. oneidensis*, functional parallels between CccR and SoFic remain enigmatic.

The few available crystal structures of complexes between Fic enzymes and their target proteins have been obtained with [34, 49] and without covalent complex crosslinking [23, 25]. In case of non-covalent complexes, including our structure, the catalytic histidine of the Fic enzyme is substituted with alanine and the target protein is AMPylated, trapping the interaction partners in a non-catalytic and therefore stabilized deAMPylation complex [25, 47]. Notably, our SoFic:EF-Tu complex crystal structure is, to our knowledge, the first report of a bacterial Fic enzyme interacting with a bacterial target (Fig. 5).

An important interface between SoFic and EF-Tu is formed by the β-flap of SoFic (Fig. 5B, Fig. S12), a conserved structural feature of Fic enzymes that is commonly assumed to confer target recognition [24, 58]. Initially, its role was revealed in the complex between IbpA’s Fic domain and Cdc42 (PDB 4ITR [23]), where the β-flap of the Fic domain forms β-sheet-like interaction to stabilize switch-I of Cdc42. In analogy to our complex, switch-I of Cdc42 is displaced in this structure, to allow accommodation of the modified residue in the active site [23, 58]. Also, the interface between the two human proteins FicD and BiP includes an intermolecular β-sheet between the β-flap of FicD and the AMPylated loop of BiP (PDB 7B7Z, 7B80 [25]). The fact that this interaction is not observed in the covalently linked FicD:BiP complex (PDB 6ZMD [49]) may correlate with its catalytic state: while the non-covalent complexes are captured in a deAMPylation state, the covalent linkage with an ATP analog is likely to result in a post-AMPylation state [49, 25]. After successful catalysis, conformational changes may induce dissociation of the complex, which can explain the lack of interactions in the post-catalytic crystal structure. We therefore conclude that the target recognition by the β-flap of Fic enzymes is universal for Fic enzymes and target proteins of both bacterial and eukaryotic origin.

Taken together, this study identified EF-Tu as a novel AMPylation target of the *S. oneidensis* Fic enzyme SoFic and provides insights into the enzyme:target interaction that corroborate previous findings in the target recognition mechanism of Fic enzymes. The analysis of the functional regulation and DNA-binding capabilities of SoFic provides a valuable basis for future studies dissecting the physiological role of SoFic for *Shewanella oneidensis*, potentially including a dual activity as transcriptional and translational regulator by promotor DNA binding and EF-Tu AMPylation.

## Materials and Methods

### Bacterial strains and genomic DNA

Recombinant protein expression was performed in *E. coli* BL21(DE3) (NEB or Promega). *Shewanella oneidensis* MR-1 (ATCC 700550 [30]) was kindly provided by the group of Professor Rainer Krull, TU Braunschweig, Germany. The genomic DNA (gDNA) of both strains was purified using the Monarch® Spin gDNA Extraction Kit (New England Biolabs, NEB) according to the manufacturer’s instructions.

### Database research to identify the AMPylation target in *E. coli*

After an AMPylation signal at a molecular weight of 45 kDa was observed during SoFic expression in *E. coli*, the genome of the K12 strain (proteome ID: UP000000625, taxon ID: 83333) was searched for proteins with a molecular weight of 40-50 kDa. The resulting 581 proteins were BLASTed (NCBI) against the *S. oneidensis* MR-1 genome (proteome ID: UP000008186, taxon ID:211586). We identified four *E. coli* proteins with sequence identity above 85 %, including EF-Tu1 (Uniprot ID: P0CE47), EF-Tu2 (Uniprot ID: P0CE48), Rho (Uniprot ID: P0AG30) and MetK (Uniprot ID: P0A817). As only a single, C-terminal amino acid varies between EF-Tu1 and EF-Tu2 in E. coli, we chose EF-Tu1 for all further studies. Furthermore, FtsZ (Uniprot ID: P0A9A6) was included based on previous research on the Fic enzyme CccR [15], which structurally resembles SoFic.

### Genes and DNA fragments

The gene sequences for SoFic 1-372 (KEGG son:SO_4266, Uniprot ID: Q8E9K5) and *S. oneidensis* EF-Tu2 (KEGG son:SO_0229, Uniprot ID: Q8EK70) and all primers used for molecular cloning were ordered from Integrated DNA Technologies (IDT). The *E. coli* genes mentioned above and EF-Ts (Uniprot ID: P0A6P1) were amplified from the *E. coli* gDNA by PCR. *S. oneidensis* gDNA was used as a PCR template to amplify a 300 bp long DNA fragment (position 4443980-4444279 in the reference genome, NCBI Accession: GCF_000146165.2) that is located upstream of the SoFic gene *mloA* (complement to position 4442861-4443979) on the *S. oneidensis* genome. This fragment was further used as a PCR template to amplify three non-overlapping 100 bp fragments that collectively span the entire original sequence. Thereby, the 300 bp sequence was effectively split into three segments of 100 bp, named F1, F2 and F3 from closest to furthest distance from the ORF. A randomized 100 bp long DNA fragment was ordered as single stranded oligos (IDT). Hybridization was performed as described before [7]. Briefly, the single stranded forward and reverse strands were mixed in a 1:1 molar ratio and heated to 95 °C for 5 min, followed by slow cooling to room temperature. To remove impurities from synthesis, the dsDNA was subsequently separated via agarose gel electrophoresis and purified from gel using a NucleoSpin Gel and PCR Clean-up Kit (MACHEREY-NAGEL, MN).

### Agarose gel electrophoresis

To verify the DNA fragments used for DNA binding by SoFic, 100 ng DNA were supplemented with 6x Gel Loading Dye, Purple (NEB) prior to electrophoresis. Agarose gels (3 % *w/v*) were prepared using 0.5 x concentrated TRIS-Borat-EDTA (TBE) buffer and supplemented with GelStar™ Nucleic Acid Gel Stain (Lonza) at a ratio of 1:50000. The electrophoresis was performed for 2 h at 50 V (5 V/cm gel) using 0.5 x TBE as a running buffer.

### Plasmid construction

SoFic constructs for protein purification were expressed using a modified pSF expression vector (Oxford Genetics) containing an N-terminal His_10_-GFP-tag and full-length SoFic. Initially, the plasmid contained a Strep-tag®II (IBA Lifesciences) and an enterokinase cleavage site between GFP and SoFic. This tag is present in the protein used to generate the complex crystal structure between SoFic^H198A^ and EF-Tu^T62A^_AMP_ but was removed from the plasmid prior to the recombinant expression of all proteins used in biochemical assays. The SoFic gene was furthermore cloned into a modified pET28b backbone (Novagen) downstream of a GFP-tag for (co-) expression experiments. EF-Ts, EF-Tu and other putative AMPylation targets were expressed using a pMAL vector (NEB) containing a His_6_-MBP-tag upstream of the gene of interest. The pSF and pMAL vectors encode for ampicillin resistance, pET28b includes a kanamycin resistance gene. All plasmids contain a tobacco etch virus (TEV) protease cleavage site between the tag and the protein of interest. For plasmid construction, vector and insert were amplified using overlapping primers and Q5® High-Fidelity DNA Polymerase (NEB), separated by agarose gel electrophoresis and purified with the Monarch® DNA Gel Extraction Kit (NEB). The fragments were ligated by sequence- and ligation-independent cloning (SLIC)[59] using T4 DNA Polymerase (NEB). Site directed mutagenesis was performed to introduce point mutations using the Q5® Site-Directed Mutagenesis Kit (NEB) according to the manufacturer’s instructions. After transformation into One Shot™ Mach1™ cells (Invitrogen), cultivation and subsequent plasmid purification with the PureYield™ Plasmid Miniprep System (Promega), all constructs were confirmed by Sanger Sequencing (Microsynth AG).

### LC-MS-based bottom-up proteomics of SDS-gel bands

#### In-gel tryptic digest

In-gel digestion was performed based on the protocol from Shevchenko et al. [60]. Shrinking and swelling was performed with 100 % (v/v) ACN and 100 mM NH_4_HCO_3_. In-gel reduction was achieved with 10 mM dithiothreitol (dissolved in 100 mM NH_4_HCO_3_). Alkylation was performed with 55 mM iodacetamide (dissolved in 100 mM NH_4_HCO_3_). Proteins in the gel pieces were digested by covering them with a trypsin solution (8 ng/µL sequencing-grade trypsin, dissolved in 50 mM NH_4_HCO_3_) and incubating the mixture at 37 °C overnight. Tryptic peptides were yielded by extraction with 2 % (v/v) FA, 80 % (v/v) ACN. The extract was evaporated. For LC-MS/MS analysis, samples were dissolved in 20 µL 0.1% (v/v) FA.

#### LC-MS/MS analysis

Chromatographic separation of peptides was achieved with a two-buffer system (buffer A: 0.1 % (v/v) FA in H_2_O, buffer B: 0.1 % (v/v) FA in ACN) on a nano-UHPLC (Dionex Ultimate 3000 UHPLC system, Thermo Fisher). Attached to the UHPLC was a peptide trap (100 µm x 20 mm, 100 Å pore size, 5 µm particle size, C18, Nano Viper, Thermo Fisher) for online desalting and purification, followed by a 25 cm C18 reversed-phase column (75 µm x 250 mm, 130 Å pore size, 1.7 µm particle size, peptide BEH C18, nanoEase, Waters). Peptides were separated using an 80 min method with linearly increasing ACN concentration from 2 % (v/v) to 30 % (v/v) ACN over 60 min.

MS/MS measurements were performed on a quadrupole-ion-trap-orbitrap MS (Orbitrap Fusion, Thermo Fisher). Eluting peptides were ionized using a nano-electrospray ionization source (nano-ESI) with a spray voltage of 1,800 V and analyzed in data dependent acquisition (DDA) mode. For each MS1 scan, ions were accumulated for a maximum of 120 ms or until a charge density of 2 × 10^5^ ions (AGC Target) was reached. Fourier-transformation based mass analysis of the data from the orbitrap mass analyzer was performed covering a mass range of m/z 200 – 1,300 with a resolution of 120,000 at m/z = 200. Peptides with charge states between 2+ - 5+ above an intensity threshold of 1,000 were isolated within a m/z 1.6 isolation window in Top Speed mode for 3 s from each precursor scan and fragmented with EThcD at 50 ms ETD reaction time and 15 % supplemental activation. MS2 scanning was performed using an ion trap mass analyzer at a rapid scan rate, covering a mass range starting at m/z 120 and accumulated for 118 ms or to an AGC target of 1 × 10^4^. Already fragmented peptides were excluded for 30 s.

#### Data processing

Raw data was searched in Proteome Discoverer (Version 3.1.0.638) with the Sequest algorithm against a reviewed *Escherichia coli* (strain B / BL21-DE3) database (Taxonomy: 469008) obtained in August 2024 containing 4,173 entries. Carbamidomethylation was set as a fixed modification for C residues. The oxidation of M, the acetylation of the protein N-terminus and the AMPylation (Phosphoadenosine) of H, K, S, T and Y were allowed as variable modifications. A maximum number of two missing tryptic cleavages was set. Peptides between 6 and 144 amino acids were considered. A strict cutoff (FDR < 0.01) was set for peptide and protein identification. The ptmRS node was used to calculate the modification site probabilities.

The mass spectrometry proteomics data have been deposited to the ProteomeXchange Consortium via the PRIDE[61] partner repository with the dataset identifier PXD071331.

### Enzyme and target co-expression experiments

For the co-expression of two proteins, 100 ng of each plasmid were transformed into electrocompetent *E. coli* BL21(DE3) cells by electroporation. After rescue for 1 h in Super Optimal Growth (SOC) medium at 37 °C and 350 rpm, the cultures were plated onto antibiotic containing LB-Agar plates and incubated at 37 °C overnight. Multiple colonies were used to inoculate 10 mL LB medium supplemented with appropriate antibiotics and grown until saturation (6-8 h) at 37 °C and 160-180 rpm. Subsequently, the cultures were diluted 1:50 into the wells of a sterile 24-well plate with lid (Sarstedt) with a final volume of 1 mL culture per well. A Spark plate reader (Tecan) was used to incubate the plates at 37 °C and 120 rpm orbital shaking (amplitude 5 mm). In intervals of 15 min, shaking was paused to measure absorbance at 600 nm and GFP fluorescence (λ_Ex_=460 nm, λ_Em_=535 nm, monochromator, bandwith 10 nm). At an absorbance value corresponding to an optical density at 600 nm wavelength (OD_600nm_) of 0.6 in a reference well, 1 mM isopropyl-β-D-thiogalactopyranosid (IPTG) final concentration were injected into specified wells, and the temperature was reduced to 22 °C. Usually, two wells were prepared per condition to compare uninduced with induced cultures. After 16 h of cultivation, a culture sample normalized by the final absorbance value was taken from each condition and centrifuged at 16,000 *g* for 1 min. Cell pellets were subsequently dissolved in homemade Laemmli Buffer (50 mM Tris-HCl, pH 6.8, 2 % (w/v) SDS, 10 % (v/v) glycerol, 100 mM dithiothreitol (DTT), 0.001 % (w/v) bromophenol blue) and boiled at 95 °C for 5 min, before separation in sodium dodecyl-sulphate polyacrylamide gel electrophoresis (SDS-PAGE), followed by Western blotting as described below.

### Recombinant protein expression for purification

Chemically competent *E. coli* BL21(DE3) cells were transformed with 100 ng plasmid DNA by heat shock, followed by rescue and plating as described above. Here, whole plates were harvested into 40 mL LB medium supplemented with appropriate antibiotics and grown for ca. 4 h at 37 °C and 160-180 rpm. These pre-cultures were diluted 1:25 into fresh, antibiotic containing LB medium in Erlenmeyer flasks and grown at 37 °C and 180 rpm until an OD_600nm_ of 0.5-0.7 was reached. Protein expression was induced with 0.5 mM IPTG and the temperature was reduced to 22 °C for overnight expression. The cultures were harvested at 6,000 g for 15 min, washed with phosphate buffered saline (PBS, 137 mM NaCl, 2.7 mM KCl, 10 mM Na_2_HPO_4_, 1.8 mM KH_2_PO_4_), centrifuged again and stored at −20 °C until purification.

### Protein Purification

#### Cell lysis

Generally, bacterial cell pellets were thawed on ice and resuspended in 50-100 mL cold Buffer A (composition for each protein described below) per 1 L expression culture. Following incubation with a spatula tip of DNaseI (AppliChem) for 10 min on ice, cells were lysed with a Constant Cell Disruption Unit (Constant Systems) at 1.8 kbar. The lysates were immediately supplemented with 1 mM phenylmethanesulfonyl fluoride (PMSF) and 40 mM (SoFic) or 25 mM (EF-Tu) imidazole were added. The lysates were cleared by centrifugation at 50,000 g at 4 °C for at least 30 min.

#### Purification of SoFic

Cleared lysates were passed over a 5 mL EconoFit Nuvia^TM^ immobilized metal affinity chromatography (IMAC) Column (Bio-Rad) equilibrated in Buffer A (50 mM HEPES-NaOH pH 8, 500 mM NaCl, 1 mM MgCl_2_, 1 mM Tris(2-carboxyethyl)phosphine (TCEP)) supplemented with 40 mM imidazole. The column was washed with 10 column volumes (C*V*) of the same buffer, followed by elution of the fusion proteins in a gradient from 30-350 mM imidazole in Buffer A over 10 *CV*. The fractions of interest were pooled and mixed with in-house prepared TEV protease in a mass ratio of 1:75 and dialyzed against Dialysis Buffer (20 mM HEPES-NaOH pH 8, 150 mM NaCl, 1 mM MgCl_2_, 0.5 mM TCEP) at 4 °C overnight. Untagged SoFic was separated from the protease, His_10_-GFP-tag, and remaining fusion proteins via reverse IMAC: the dialyzed sample was passed over a column that was equilibrated and washed in Buffer A. Untagged SoFic proteins bound to the IMAC column unspecifically and were eluted with 25 mM imidazole in Buffer A. Fractions of interest were pooled and concentrated to a volume ≤ 2 mL using Amicon® Ultra Centrifugal Filter units (Merck Millipore). After centrifugation for 5 min at 21,000 *g* and 4 °C, the sample was injected onto a HiLoad 16/600 Superdex 75 pg column (Cytiva) equilibrated in Buffer C (20 mM HEPES-NaOH pH 8, 150 mM NaCl, 1 mM MgCl_2_, 1 mM DTT) for size exclusion chromatography (SEC). Elution peaks were analyzed via SDS-PAGE and the fractions of highest purity were pooled, concentrated using Amicon® Ultra Centrifugal Filter units and stored at −80 to −75 °C after freezing in liquid nitrogen. The molar concentration of the samples was determined via UV-Vis spectroscopy at 280 nm (ε = 51,340 M^−1^cm^−1^).

#### Small Scale Purification of SoFic

For the purification of small amounts His_10_-GFP-tagged SoFic mutants, cell pellets of 100-300 mL expression cultures were resuspended in 10 µL per mg pellet xTractor™ Buffer (Takara Bio) supplemented with 1 mM PMSF and a spatula tip of DNaseI and incubated under agitation at 4 °C for 1 h. After centrifugation as described above, the cleared lysates were passed over 250 µL bed volume Protino Ni-NTA Agarose beads (MN) twice by gravity flow. The beads were equilibrated in Buffer A substituted with 40 mM imidazole prior to loading the lysates and washed with the same buffer afterwards. The proteins were eluted using Buffer A supplemented with 150 mM imidazole in five fractions of 0.5 µL. Fractions of interest were pooled and dialyzed into Buffer C overnight. The molar concentration of the fusion proteins was determined via UV-Vis spectroscopy at 488 nm (GFP-fluorescence, ε =55,900 M^−1^cm^−1^).

#### Purification of EF-Tu and EF-Ts

Cleared lysates containing His_6_-MBP-tagged EF-Tu were passed over a 5 mL EconoFit Nuvia IMAC Column, which was equilibrated and washed with EF-Tu Buffer A (50 mM HEPES-NaOH pH 7.5, 500 mM NaCl, 1 mM MgCl_2_, 1 mM TCEP, 5 % (v/v) glycerol, 10 µM guanosine diphosphate (GDP)) supplemented with 25 mM imidazole. Elution was performed over 10 *CV* in a gradient from 25-350 mM imidazole EF-Tu Buffer A. Protein containing fractions were pooled, mixed with in-house prepared TEV protease in a mass ratio of 1:75 and dialyzed against EF-Tu Dialysis Buffer (20 mM HEPES-NaOH pH 7.5, 150 mM NaCl, 0.5 mM TCEP, 5 % (v/v) glycerol, 10 µM GDP) at 4 °C overnight. The dialyzed sample was passed over a 5 mL MBPTrap™ HP column (Cytiva) coupled to an IMAC column to completely capture the 6xHis-MBP-tag. Untagged EF-Tu bound to the IMAC column unspecifically and was eluted with 25 mM imidazole in Buffer A after washing with Buffer A and removal of the MBP-trap. EF-Tu containing fractions were concentrated and purified by SEC as described above in EF-Tu Buffer C (20 mM HEPES-NaOH pH 7.5, 150 mM NaCl, 1 mM DTT, 5 % (v/v) glycerol, 10 µM GDP). Elution peaks were analyzed via SDS-PAGE and the fractions of highest purity were pooled, concentrated and stored as above. The molar concentration of the samples was determined via UV-Vis spectroscopy at 280 nm (ε = 20,400 M^−1^cm^−1^ (EF-Tu) + 6,840 M^−1^cm^−1^ (GDP) = 27,240 M^−1^cm^−1^ to account for the absorption of GDP bound by EF-Tu). EF-Ts was purified identical to EF-Tu using buffers without glycerol and GDP. The molar concentration of EF-Ts was determined via UV-Vis spectroscopy at 280 nm as well (ε = 4,470 M^−1^cm^−1^). The concentrated protein samples were stored at −80 to −75 °C after freezing in liquid nitrogen.

To remove GDP from EF-Tu, previously purified protein was incubated with 10 mM EDTA in nucleotide free EF-Tu Buffer C (20 mM Hepes (pH 7.5), 150 mM NaCl, 5 % (v/v) glycerol, 1 mM DTT, 5 mM EDTA) for 1 h at 37 °C. It was then subjected to SEC in the same buffer to separate protein from unbound nucleotides. The peak fractions were concentrated, followed by removal of EDTA by SEC in 20 mM Hepes (pH 7.5), 150 mM NaCl, 5 % glycerol, 1 mM DTT. The molar concentration of the concentrated peak fractions was determined via UV-Vis spectroscopy at 280 nm (ε = 20,400 M^−1^cm^−1^) and the samples stored as described above.

### Preparative AMPylation of EF-Tu

EF-Tu was preparatively AMPylated after removal of the His-MBP-tag during purification. At this stage, the protein was present in EF-Tu Buffer A containing 25 mM imidazole. His_10_-GFP-tagged SoFic^E73G^ was added to a molar ratio of 1:20. The reaction was started by the addition of 1.5 mM ATP and 5 mM MgCl_2_ and incubated at 20 °C overnight. The solution was subsequently concentrated and subjected to SEC as described for unmodified EF-Tu to remove the GFP-SoFic fusion protein. The molar concentration of the concentrated samples after SEC was determined via UV-Vis spectroscopy at 280 nm (ε = 20,400 M^−1^cm^−1^ (EF-Tu) + 6,840 M^−1^cm^−1^ (GDP) + 3,560 (AMP) = 30,800 M^−1^cm^−1^ to account for the absorption of GDP bound by EF-Tu and the covalently attached AMP). Intact LC-MS analysis was used to confirm ≥ 98 % modification.

### Nucleotide exchange of EF-Tu

To load EF-Tu with mant-labeled GDP or GppNHp, 10 nmol purified protein in the GDP-bound state were mixed with 100 nmol (10x excess) of mant-labeled nucleotide (Jena Bioscience) in nucleotide-free EF-Tu Buffer C (20 mM HEPES-NaOH pH 7.5, 150 mM NaCl, 1 mM DTT, 5 % (v/v) glycerol). After incubation at 37 °C for 30 min, excess unbound nucleotide was removed using NAP-5 columns equilibrated in nucleotide-free EF-Tu Buffer C. Protein containing fractions were pooled and concentrated, successful nucleotide exchange was confirmed using UV-Vis spectroscopy at 355 nm to detect mant-fluorescence (ε = 5.7 mM^−1^cm^−1^). For storage at −80 °C, the protein solutions were supplemented with 10 µM of the respective nucleotide.

### Complex formation of EF-Tu and EF-Ts

Prior to complex formation, 50 µM purified *ec*EF-Tu^WT^:GDP were incubated with 10 mM EDTA in nucleotide-free EF-Tu Buffer C to reduce the affinity to GDP. After incubation at 37 °C for 30 min, 50 µM purified EF-Ts were added to the reaction, followed by incubation at 37 °C for 30 min. The sample was then subjected to SEC in nucleotide-free EF-Tu Buffer C as described above. EF-Tu:EF-Ts complexes eluted in a symmetric peak and co-elution was confirmed via SDS-PAGE analysis. The peak fractions were concentrated as described above and the molar concentration of the final samples was determined via UV-Vis spectroscopy at 280 nm (ε = 24,870 M^−1^cm^−1^ for EF-Tu:EF-Ts or ε = 28,431 M^−1^cm^−1^ EF-Tu_AMP_:EF-Ts). The samples were stored as described above.

### Fluorescent labeling of SoFic

To lower the detection limit in aSEC, SoFic was labeled non-specifically with the cysteine-reactive maleimide-coupled fluorescent dyes ATTO-488 or ATTO-532 (ATTO-TEC). The lyophilized dyes were prepared and stored as described before [47], the labeling procedure was adapted as follows: the labeling steps were performed in degassed FL-Buffer (50 mM HEPES-NaOH pH 8, 150 mM NaCl, 1 mM MgCl_2_, 0.05 mM DTT). After reduction of 100 nmol purified SoFic with 10 mM DTT in a total volume of 2 mL for 10-20 min at room temperature (RT), the DTT was removed by washing four times with 15 mL FL-Buffer using an Amicon® Ultra Centrifugal Filter units. Concentration of the washed sample was determined and the dye was added to a molar ratio of 1.1:1 (dye:protein). After light-protected incubation overnight at 5-10 °C, the unbound dye was removed by washing the sample seven times as described above. The sample concentration and degree of labeling (DOL) were determined using UV-Vis spectroscopy at 280 nm for the protein concentration and 488 nm for ATTO-488 or 532 nm for ATTO-532 concentrations. The DOL was calculated as a ratio of dye concentration to protein concentration, the described labeling procedure generally resulted in 95-100 % labeling efficiency (indicating ca. 1 molecule dye per molecule SoFic).

### Intact liquid chromatography mass spectrometry (LC-MS)

LC-MS was used to verify purified protein samples and to analyze the ratio of AMPylated to unmodified protein species. In all cases, protein samples were diluted to 0.05 mg/mL in LC-MS-grade water (Chemsolute®) and centrifuged for 5 min at 21000 *g*, 4 °C. A volume of 2 µL per sample were injected onto a ProSwift™ RP-4H 1×50 mm column (Thermo Fisher Scientific) for desalting. Elution was performed with an acetonitrile gradient, followed by injection into a maXis II ETD ESI LCMS (Bruker Daltonics). Mass spectra were deconvoluted and analyzed using DataAnalysis (Version 5.1, Bruker Daltonics). The expected MW for each protein was calculated based on the amino acid sequence using ProtParam [62].

### Nano differential scanning fluorimetry

Purified EF-Tu samples were diluted to 0.15 mg/mL in EF-Tu Buffer C. All samples were centrifuged at 20,000 *g* for 5 min at 4 °C before loading into Prometheus Standard Capillaries (NanoTemper Technologies). Protein fluorescence at 350 nm (*F*_350_) and 330 nm (*F*_330_) was measured over a temperature range from 15-80 °C using a Prometheus nDSF (NanoTemper Technologies) device. Data was analyzed with the MoltenProt Webserver[63, 64]. The melting temperatures (*T*_m_) were obtained from the inflection points of the *F*_350_/*F*_330_-ratio vs. temperature plots and averaged over at least three technical replicates each for two independently purified protein batches.

### AMPylation activity assays

For all assays, the concentration of EF-Tu was 10 µM (0.43 mg/mL), SoFic was added to 0.1 µM (molar ratio of 1:100). The reactions were performed in Buffer C (20 mM HEPES-NaOH pH 8, 150 mM NaCl, 1 mM MgCl_2_, 1 mM DTT) and started by the addition of 1.5 mM ATP, followed by incubation at 15 °C for 30-240 min (AMPylation kinetics) or overnight (endpoint measurements). For analysis by intact LC-MS, the reactions were diluted to 0.05 mg/mL EF-Tu in LC-MS-grade water. For Western blot analysis, the reactions were stopped by the addition of Laemmli buffer and boiling at 95 °C for 5 min. A quantity of 250 ng of EF-Tu was applied to SDS-PAGE. Densitometric quantification of the AMPylation kinetics was performed using ImageJ [65]. After background subtraction for each anti-AMP signal, the percentage of AMPylated EF-Tu was calculated relative to the anti-AMP signal of the positive control (100 % AMPylated EF-Tu^T62A^). We did not normalize the relative signals to the loading controls, because high levels of AMPylation randomly interfered with silver staining of EF-Tu (see e.g., Fig. S7B, lane 7, Fig. S7C, lane 2).

### deAMPylation activity assays

The deAMPylation was analyzed by incubation of 10 µM AMPylated EF-Tu^T62A^ with 0.1 µM SoFic. The reactions were prepared in Buffer C and incubated at 15 °C for 5-40 min. The reactions were stopped by the addition of Laemmli buffer and boiling at 95 °C for 5 min. A quantity of 250 ng of EF-Tu was applied to SDS-PAGE. Densitometric quantification of the deAMPylation kinetics was performed as described above. Here, the percentage of AMPylated EF-Tu was calculated relative to the anti-AMP signal of the negative control (reaction without SoFic, t = 40 min).

### Western blotting and antibody detection

After separation on 12 % (w/v) polyacrylamide gels (homemade), protein or bacterial culture samples were transferred on Immobilon®-P PVDF membranes (Merck-Millipore) for 1.5 h at 230 mA using a Biometra Fastblot B44 (Analytik Jena) device. The membranes were blocked with 1x Roti®-Block (Carl Roth) in Tris-buffered saline with 0.1 % (v/v) Tween20 (TBS-T, homemade) for at least 1 h at RT. For detection of AMPylated proteins, a mouse-anti-AMP monoclonal antibody 17G6 [39] was added to the blocking solution at a concentration of 0.5 µg/mL, followed by incubation at 4 °C overnight. The membranes were washed three times for 10 min in TBS-T before incubation for 1 h at RT with the secondary antibody goat anti-mouse IgG (H+L) HRP conjugate (#31430, Thermo Fisher Scientific) at a ratio of 1:20000 in TBS-T. The membrane was washed again as above and incubated for 5 min at RT with a 1:1:1:1 mixture of the SuperSignal West™ reagents Pico PLUS and Dura (Thermo Fisher Scientific) to develop the peroxidase signal. Chemiluminescence was detected with an Intas ECL Chemocam (Intas Science Imaging Instruments).

Prior to the repeated probing of the same membrane using alternative antibodies, stripping of the Western blots was performed using Roti^®^Free Stripping-Buffer 2.0 (Carl Roth). The solution was pre-warmed to 37 °C, added to the membrane for incubation at RT for 1 h. After washing five times for 10 min in TBS-T, the blocking and detection procedures described above were repeated.

To detect GFP-tagged SoFic constructs in bacterial culture samples, we used the rabbit GFP polyclonal antibody (Invitrogen, #A-11122) at a ratio of 1:2,000 in 1x Roti®-Block in TBS-T and the goat anti-rabbit IgG H&L (HRP) preadsorbed (#ab7090, abcam) at a ratio of 1:20,000 in TBS-T as a secondary antibody. His6-MBP-EF-Tu fusion proteins were detected using the HisProbe™-HRP Conjugate (Thermo Scientific™, #15165) at a ratio of 1:5,000 in TBS-T.

To visualize untagged EF-Tu in in vitro assays, the membrane was stripped as described above and subsequently stained for 10 min at RT using a colloidal silver solution (0.8 % (w/v) ferrous sulfate heptahydrate, 2 % (w/v) sodium citrate dihydrate, 0.2 % (w/v) silver nitrate). The stained membrane was washed thoroughly in ddH2O before imaging.

### Stopped-flow kinetics

Binding kinetics for GDP, GppNHp and EF-Ts were recorded using a SX20 stopped-flow spectrophotometer (AppliedPhotophysics) as described previously [46]. All experiments were performed at 25 °C and in degassed buffer composed of 20 mM Hepes pH 7.5, 150 mM NaCl, 1 mM MgCl_2_, 1 mM DTT. Equal volumes of the reagents were rapidly mixed, all concentrations given below refer to the stock solutions before mixing. Nucleotide binding was monitored via FRET between the single Trp (W178) of EF-Tu and mant-labeled nucleotides. When bound to EF-Tu, mant-fluorescence can be indirectly induced by Trp excitation at 280 nm and detected after passing a 400 nm cutoff filter. Thus, association or dissociation of the labeled nucleotide to EF-Tu leads to an increase or decreases of mant-fluorescence, respectively. The nucleotide exchange reactions were monitored over time as specified below. For dissociation kinetics, 0.5 µM EF-Tu bound to mant-GDP or mant-GppNHp were rapidly mixed with 50 µM unlabeled GDP (1500 s detection) or GTP (500 s detection). The k_off_ [s^−1^] rates were directly obtained by fitting the resulting curves to an exponential function:

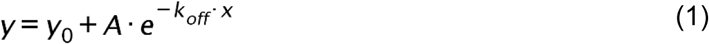

The k_off_ rates were averaged across 3-5 measurements for each condition. To record association kinetics, 0.5 µM nucleotide-free EF-Tu were mixed with different concentrations (0; 2; 4; 6; 10; 12 µM) mant-GDP (2 s detection) or mant-GppNHp (5-20 s detection). Exponential fitting of the fluorescence signal curves to the exponential function

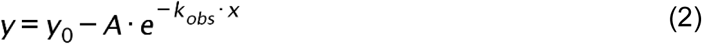

yielded concentration dependent k_obs_ [s^−1^] rates. The k_obs_ rates were averaged across 3-5 measurements for each condition before plotting over nucleotide concentration (*c*). The k_on_ [µM^−1^s^−1^] rates were then obtained as the slope of a linear fit to the individual data points:

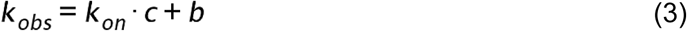

K_D_ was calculated as the ratio:

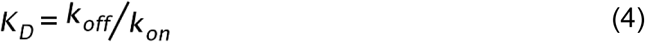

Dissociation kinetics of EF-Ts were monitored by the change in Trp fluorescence of EF-Tu, since no Trp residue is present in EF-Ts. Nucleotide free EF-Tu:EF-Ts complex at 1 µM was mixed with 300 µM GDP or GTP to induce complex dissociation (0.05 s detection). The k_off_ [s^−1^] rates were directly obtained by fitting the resulting curves to an exponential function as above (Equation (1)). All data analysis was performed in OriginPro, Version 2023 (OriginLab Corporation).

### In vitro expression assay

Coupled *in vitro* transcription and translation reactions were prepared using an *E. coli* S30 Extract System for Circular DNA (Promega). Amino acids, S30 premix and S30 extract were combined in ratios according to the manufacturer’s instructions in a total reaction volume of 20 µL. The reactions were supplemented with 0.05-10 µM EF-Tu^WT^ or EF-Tu^WT^_AMP_. EF-Tu Buffer C was used as a control. The reactions were started by the addition of 780 ng of a GFP encoding plasmid (T5 promotor) in the wells of a transparent non-binding flat bottom 384 well plate (Corning Incorporated). The plate was incubated in a Spark plate reader (Tecan) at 37 °C and GFP fluorescence (λ_Ex_=485 nm, λ_Em_=535 nm, bandwidth 20 nm, monochromator) was monitored in intervals of 2 min for 2 h.

### Analytical size exclusion chromatography

Analytical SEC was performed using a Prominence high pressure liquid chromatography (HPLC) system (Shimadzu). All protein samples were supplemented with 12 µM Vitamin B12 (Sigma Aldrich) as an internal standard and 90 µL were injected onto a Superdex^TM^ 75 pg Increase column (10/300 GL, Cytiva) for homodimerization analyses or a Superdex^TM^ 200 pg Increase column (10/300 GL, Cytiva) for DNA binding and protein interaction experiments. The samples were eluted with 30 mL Buffer C at a flow rate of 0.5 mL/min. Absorbance channels were set to 280 nm (A_280 nm_) for protein and 260 nm (A_260 nm_) for DNA detection. The fluorescence detector was set to λ_Ex_=280 nm, λ_Em_=340 nm to detect Trp fluorescence, λ_Ex_=475 nm, λ_Em_=550 nm for ATTO-488 and λ_Ex_=520 nm, λ_Em_=560 nm for ATTO-532 detection. Analysis of all chromatography traces was performed in OriginPro, Version 2023 (OriginLab Corporation).

For the interaction between SoFic and EF-Tu (Fig. S11), unprocessed aSEC traces are shown (accurate alignment of the internal standard in the raw data was confirmed). To compare the retention times of SoFic^WT^ and dimerization mutants (Fig. 1B), the A_280 nm_ elution profiles were baseline subtracted, normalized by their maximum intensity and aligned by the internal standard for visualization and peak analysis to extract the retention times of each protein peak. Subsequently, the retention times were converted into molecular weights (MW_calc_, Table S1) using a linear regression of the retention times obtained for a molecular weight standard (BioRad).

To calculate the apparent K_D_ for dimerization of SoFic (Fig. 1C), the chromatography traces were baseline subtracted and aligned by the internal standard peak. A_280 nm_ detection was sufficient to detect concentration above 0.09 µM. Lower concentrations were analyzed using the fluorescence traces after accounting for the delay between both detectors by subtraction of the delay time. The retention times of the protein peaks were extracted from the processed traces, converted into MWs as above and normalized from 0 % (lowest apparent MW) to 100 % (highest apparent MW) dimer (fraction dimer, *f_D_*). The resulting values were plotted over protein concentration (*c*) and fitted to a hyperbolic formula to obtain K_D_:

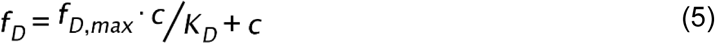

The chromatograms of DNA binding experiments (Fig. S2) were baseline subtracted before visualization (accurate alignment of the internal standard in the raw data was confirmed).

### AlphaFold2 prediction of the SoFic:EF-Tu complex structure

The SoFic:EF-Tu complex structure was predicted based on the full length sequences of SoFic (Uniprot ID: Q8E9K5) and *ec*EF-Tu (Uniprot ID: P0CE47) using the open source software ColabFold (Version 1.5.2) [41]. The prediction was run with the following parameters/settings:

- MSA mode: mmseqs2_uniref_env
- model type: alphafold2_multimer_v3
- 1 seed (= 5 models)
- 12 recycling steps
- PDB-based template search
- MSA sequence distribution: max. 508 paired and max. 2048 unpaired sequences
- Pairing strategy: greedy

Out of the five resulting structural models, the model with the lowest predicted aligned error (PAE) score was used for analysis and representation. This model also was the only one with an interface predicted template modeling (ipTM) score ≥ 0.85, which is a stringent reliability cutoff for predicted protein-protein interactions [66].

### Ray crystallography

Prior to crystallization, all proteins were diluted to 5-10 mg/mL in their respective size-exclusion chromatography buffer (Buffer C). For EF-Tu, Buffer C contained 10 µM GDP. Samples of SoFic^WT^, SoFic^L31D^ and SoFic^L50D^ were supplemented with 5 mM ATP and 5 mM MgCl_2_. Initial crystallization trials were performed using a Mosquito pipetting robot (SPT Labtech) to dispense 100 nL protein solution and 100 nL reservoir solution from the JCSG Core I–IV screens (Qiagen) as sitting drops on 96-well plates (Swissci). The crystallization conditions for the SoFic:EF-Tu complex crystal structure and all EF-Tu structures except *so*EF-Tu^T62A^_AMP_ were further optimized using manually prepared hanging drop crystallization plates, where 2 µL protein solution were mixed with 2 µL reservoir. The final individual crystallization conditions and protein concentrations are listed in Supplementary Data File 4. When the crystallization condition did not provide sufficient cryoprotection, crystals were soaked in reservoir solution supplemented with 25–30 % (v/v) glycerol before flash-cooling in liquid nitrogen using cryo-loops (Hampton Research).

Diffraction data were collected at the EMBL beamlines P13 and P14 of the PETRA III storage ring (DESY, Hamburg, Germany). All datasets were processed using XDS [67]. Structure factor amplitudes for SoFic^L31D^ and SoFic^L50D^ were scaled and merged with AIMLESS [68] via the autoPROC pipeline [69]. For datasets exhibiting significant anisotropy, an anisotropic cutoff and scaling were performed with STARANISO (Global Phasing Ltd.), using local I/σ(I) and CC_1/2_ criteria.

Initial phases were obtained by molecular replacement using PHASER [70], with the search models listed in Supplementary Data File 4. Manual model building and corrections were performed in COOT [71], and refinements were carried out using PHENIX refine [72]. For the SoFic^L31D^ structure, twin refinement was applied using the twin law *h*, *–k*, *–l*, with a refined twin fraction of 0.46.

Model quality was assessed with MolProbity [73] and the wwPDB validation server [74]. Methodological variations specific to individual structures, data collection and refinement statistics, as well as the PDB accession codes, are provided in Tables S2, S3, S5, S7 and Supplementary Data File 4.

## Data reproducibility

Expression and co-expression experiments were performed in biological triplicates. *In vitro* assays were performed in biological duplicates, i.e. using two independently expressed and purified protein samples. Stopped-flow kinetics are the only exception, where 3-5 technical replicates were performed.

## Acknowledgements

We would like to thank Dr. Dorothea Höpfner and Dr. Marietta Sandkamp-Kaspers for their advice on experimental design. A.I. acknowledges access to the core facilities and laboratories of the Centre for Structural Systems Biology Hamburg (CSSB). We acknowledge support by the Sample Preparation and Characterization (SPC) and High-throughput Crystallization (HTX) facilities at EMBL Hamburg. We would like to thank the local contacts for their assistance in using the beamlines P13 and P14 operated by EMBL Hamburg at the PETRA III storage ring (DESY, Hamburg, Germany). We thank the Core Facility Mass Spectrometric Proteomics as part of the Technology Platform Mass Spectrometry (TPMS) at University of Hamburg (UHH) and University Medical Center Hamburg-Eppendorf (UKE) for support with mass spectrometric measurements and analysis funded by the Deutsche Forschungsgemeinschaft (DFG, German Research Foundation, project no. 247377969).

## Funding

A.I. acknowledges support by the Sonderforschungsbereich (SFB) 1035 project B05 of the Deutsche Forschungsgemeinschaft (DFG, German Research Foundation, project no. 201302640). S.R. was funded by the SFB1035 and the DFG - Research Training Group 2771 (project no. 453548970). B.S. and H.S. acknowledge funding by the DFG (project no. 247377969).

## Author contributions

Design of Research: SR, AI; Protein purification and biochemical experiments: SR, AB; X-Ray crystallography and model building: VP, SR, AB; LC-MS-based proteomics: BS, HS; Alphafold prediction: MHB; Data analysis and visualization: SR; Writing-original draft: SR; Writing-review and editing: VP, AI.

## Competing Interest

Authors declare that they have no competing interests.

## Data and materials availability

All data needed to evaluate the conclusions in the paper are present in the paper and the Supplementary Materials. Structure factors and model coordinates for the crystal structures have been deposited in the RCSB PDB under the following accession codes: *ec*EF-Tu^T62A^_AMP_: 9TBU; *so*EF-Tu^WT^: 9T8G; *so*EF-Tu^T62A^: 9T9C; *so*EF-Tu^T62A^_AMP_: 9TCK; SoFic^H198A^:*ec*EF-Tu^T62A^_AMP_-complex: 9TB0; SoFic^WT^: 9TCH; SoFic^L31D^: 9TCL; SoFic^L50D^: 9TC4. The mass spectrometry proteomics data have been deposited to the ProteomeXchange Consortium via the PRIDE partner repository with the dataset identifier PXD071331. Additional data related to this paper may be requested from the authors.

## Notes

### Competing Interest Statement

The authors have declared no competing interest.

