## Supplementary Information for "The *Shewanella oneidensis* Fic enzyme SoFic targets the switch-I region of EF-Tu for AMPylation"

**This PDF includes:**

Table of Contents

Data File Captions 1-4

Figures S1-S12

Tables S1-S8

References for the SI citations

Other Supplementary Materials for this manuscript include the following:

Data Files 1-4

**Data File Captions**

**Data File 1: Database search results for the identification of the AMPylation target of SoFic.** The first sheet "*E. coli* K12 proteome 40-50 kDa" contains the hits after filtering the reference genome for *E. coli* BL21 DE3 (UniProt UP000002032, Organism 469008) for proteins with a MW of 40-50 kDa. The second sheet "BLAST Hits *S. oneidensis*" contains the hits of a BLAST search of all proteins from the first sheet against the genome of *S. oneidensis* MR-I (taxid:211586) ordered by their sequence identidy in percent (%).

**Data File 2: LC-MS/MS results for the AMPylation signal in lane 2 of Fig. 2A (SoFic$^{WT}$ -IPTG).** The sample from the Western blot was separated on another SDS-PAGE gel, stained with Coomassie and the bands at the corresponding molecular weight were excised for bottom-up proteomic analysis by LC-MS/MS.

**Data File 3: LC-MS/MS results for the AMPylation signal in lane 6 of Fig. 2A (SoFic$^{E73G}$ +IPTG).** The sample from the Western blot was separated on another SDS-PAGE gel, stained with Coomassie and the bands at the corresponding molecular weight were excised for bottom-up proteomic analysis by LC-MS/MS.

**Data File 4: Crystallization parameters and structure solution details.** This file includes information on the protein concentration, crystallization conditions, the beamline used for data collection as well as data processing details and the model used for molecular replacement (MR) for each crystal structure.

### Supplementary Figures

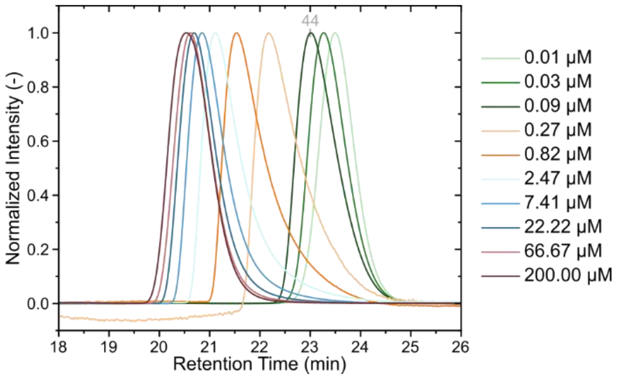

**Figure S1: The concentration dependent retention time shifts of SoFic in analytical size exclusion chromatography.**

Analytical size exclusion chromatography (aSEC) traces of SoFic at the indicated concentrations. Tryptophane fluorescence ($\lambda_{Ex}$=280 nm, $\lambda_{Em}$=340 nm) was used to detect 0.01-0.09 µM SoFic, while absorption at 280 nm was sufficient to detect all higher concentrations. Peak broadening at intermediate concentrations indicates a dynamic interconversion between monomeric and dimeric enzyme. The retention time shifts with changing concentration were used to calculate the apparent molecular weight and dimeric fraction of the enzyme to estimate the $K_D$ for dimerization. A full description of peak analysis for $K_D$ determination in Fig. 1C can be found in the method section. For this representation, all traces were baseline subtracted and normalized to the peak maximum. The retention time of the 44 kDa reference protein from a gel filtration standard (BioRad) is indicated in grey. The data is representative of two technical replicates for each of two independent biological samples.

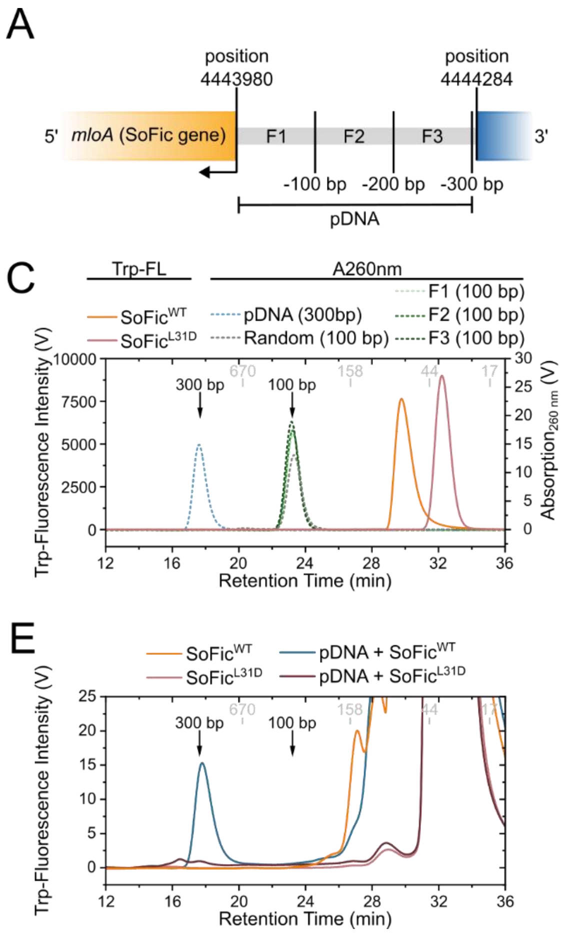

**Figure S2: SoFic binding to its own promotor DNA is sequence specific and requires dimerization**

- **A.** Schematic representation of the SoFic gene *mloA* on the complement strain of the *S. oneidensis* genome and the 3' upstream sequence separating it from the next gene. The whole 300 bp long sequence (pDNA) was further divided into three 100 bp long fragments (F1-F3) to narrow down the binding sites of SoFic.
- **B.** Agarose gel electrophoresis to validate the length of the indicated DNA analytes shown in **A**. R denotes a randomized 100 bp long control fragment with the same GC content as F1 (Table S2). 100 ng of each DNA fragment were loaded onto a 3 % (*w/v*) agarose gel in 0.5x TBE.
- **C.** Analytical size exclusion chromatography (aSEC) traces of the indicated protein and DNA analytes. The samples contained 25 µM protein and 25 nM of the 300 bp long pDNA. 75 nM were used for the 100 bp long fragments to provide equal amounts of base pairs as for the pDNA. The left intensity axis displays tryptophane (Trp) fluorescence intensity of protein samples ($\lambda_{Ex}$=280 nm, $\lambda_{Em}$=340 nm, solid lines), right intensity axis indicates the absorption at 260 nm (A260 nm) of DNA samples (dashed lines).
- **D.** Trp fluorescence intensity aSEC traces of the indicated DNA analytes in the absence of protein. The retention times of the same analytes detected via absorption at 260 nm in **C** are indicated by arrows. Since no Trp fluorescence signal is detected for the DNA analytes,

this channel is insensitive to DNA. Thus, Trp fluorescence was used to specifically detect the co-elution of protein at the retention times of DNA in mixed samples (**E,F**). Especially for the pDNA, which elutes at the void volume of the column, binding by SoFic could not be expected to cause a shift in retention time and required detection via protein specific Trp fluorescence.

**E.** Trp fluorescence intensity aSEC traces of $SoFic^{WT}$ and $SoFic^{L31D}$ in the absence and presence of the 300 bp long pDNA. The presence of a protein specific Trp fluorescence peak at the retention time of the pDNA qualitatively shows binding by dimeric $SoFic^{WT}$, but not monomeric $SoFic^{L31D}$.

**F.** Trp fluorescence intensity aSEC traces of $SoFic^{WT}$ in the absence and presence of the indicated 100 bp long DNA fragments. The intensities of the protein specific Trp fluorescence peaks at the retention time of 100 bp DNA qualitatively shows that the highest amount of protein co-elutes with F1. No co-elution is detected for the randomized 100 bp DNA fragment.

In **C-F**, black arrows indicate the retention times of the DNA analytes alone as detected in **C** and the retention times of the reference proteins from a gel filtration standard (BioRad) are indicated in grey by their molecular weights in kDa. All traces in **C-F** were baseline corrected.

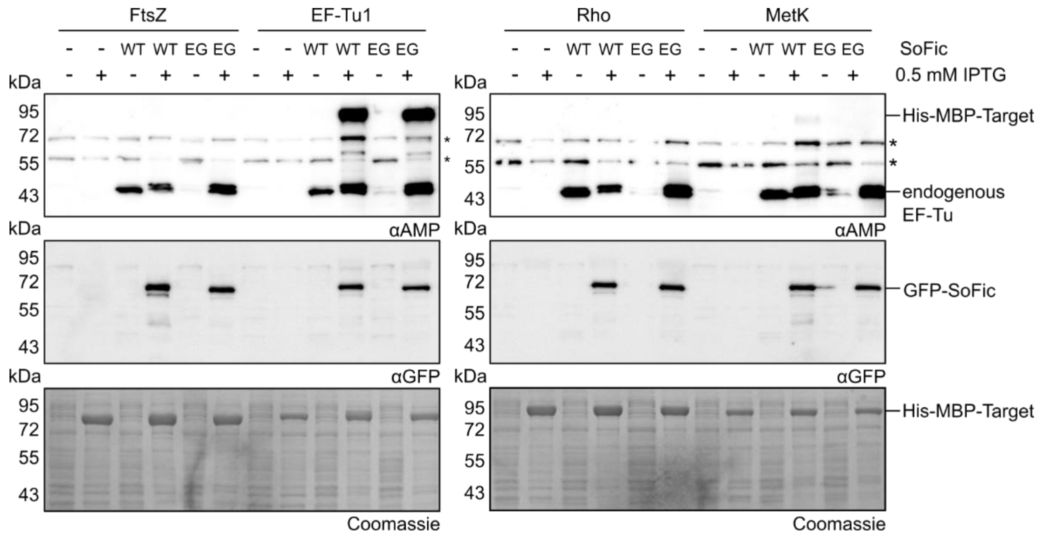

**Figure S3: Co-expression of SoFic with potential AMPylation targets reveals AMPylation of EF-Tu.**

Western blot analysis of culture samples after co-expression of GFP-tagged $SoFic^{WT}$ (WT) or $SoFic^{E73G}$ (EG) with $His_6$-MBP-tagged target candidates for AMPylation that were identified by a database research attempt (Fig. 2B). Culture samples were normalized by $OD_{600nm}$ after cultivation without or with 1 mM IPTG at 22 °C overnight for SDS-PAGE and Western blotting. After detection with an anti-AMP antibody, the membranes were stripped and detected again with an anti-GFP-antibody to control for SoFic expression. Residual protein in the SDS-PAGE gels after Western blotting was stained with Coomassie to control for overexpression of the target candidates. Asterisks (*) indicate endogenous background signals in *E. coli*.

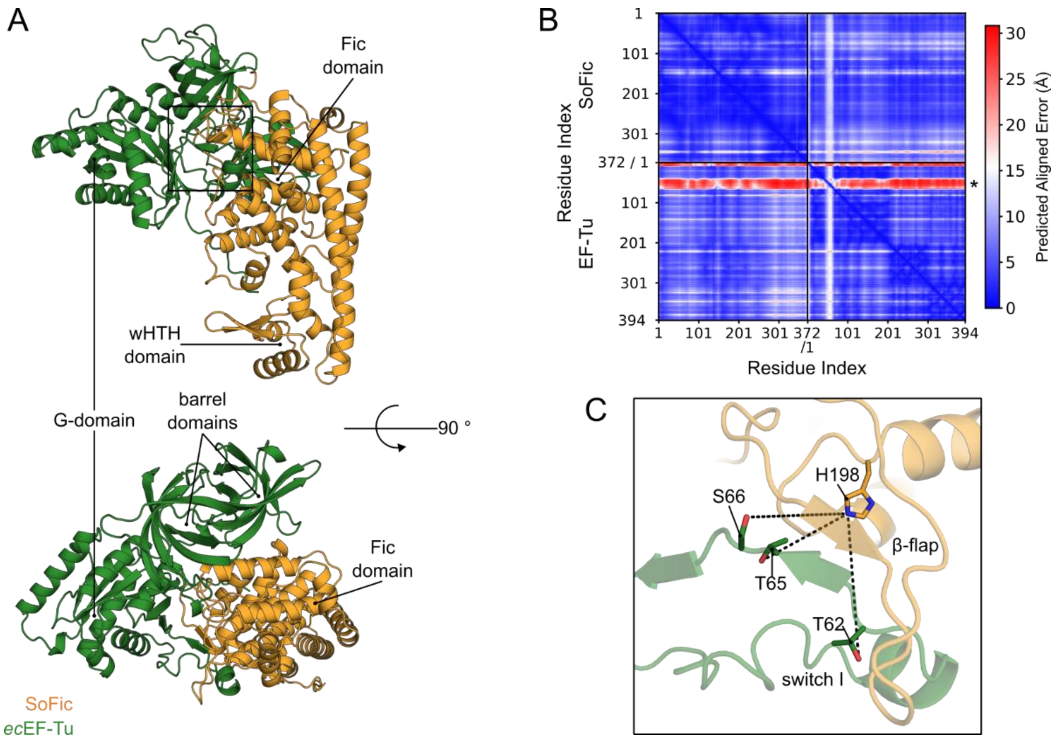

**Figure S4: The Alphafold2 prediction of a complex between SoFic and EF-Tu suggests modification sites within the switch-I region of EF-Tu.**

**A.** Front view (top panel) and top view (bottom panel) of a SoFic:EF-Tu complex structure that was predicted by Alphafold2, showing EF-Tu (green) bound to the Fic domain of SoFic (orange). One chain of each protein was used for the prediction and the interface predicted template modeling (ipTM) score for the prediction is 0.868.

**B.** Predicted aligned error plot (PAE) of the predicted complex in **A**. The red region (asterisk) is predicted with low confidence and corresponds to EF-Tu residues 39-68, including the conserved switch-I.

**C.** Close-up view of the interaction between EF-Tu and SoFic in the catalytic center of SoFic. The switch-I region of EF-Tu is rearranged into the active center, where it forms an intermolecular β-sheet with the β-flap of SoFic. Three hydroxyl group containing residues are found in close proximity to the catalytic His (H198) of SoFic: T62 (12.5 Å), T65 (9.0 Å) and S66 (12.8 Å).

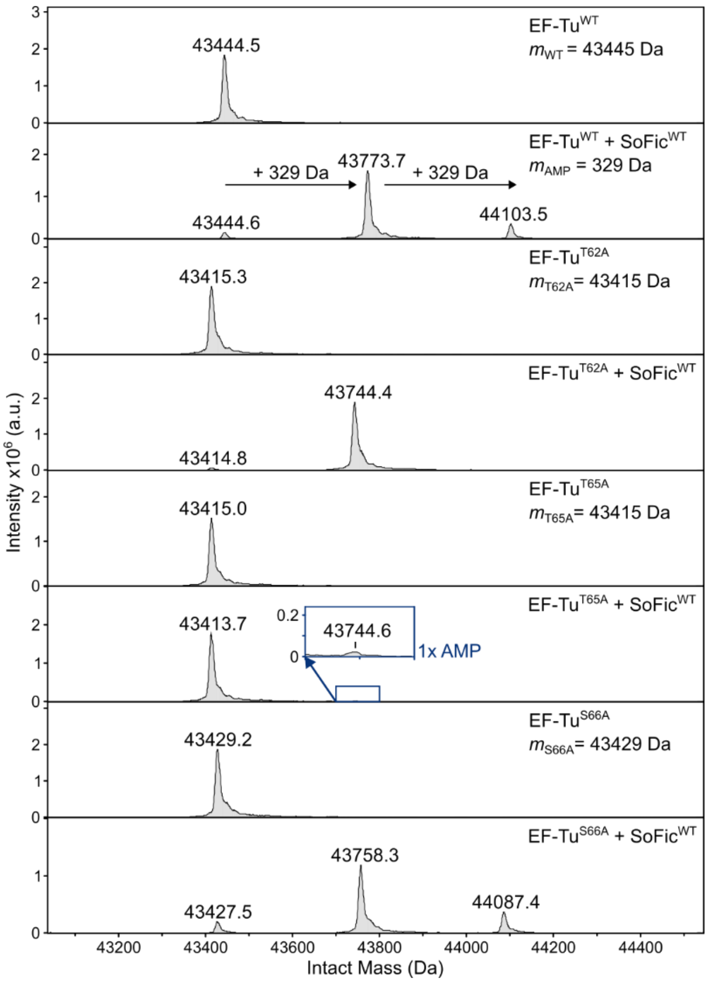

**Figure S5: EF-Tu is AMPylated at T65 as a primary and T62 as a secondary modification site.**

Intact mass spectra of the indicated EF-Tu mutants before and after incubation with 0.1 µM SoFic$^{WT}$ and 1.5 mM ATP at 15 °C overnight. The expected mass of the unmodified EF-Tu mutants is indicated, AMPylation causes a mass shift of 329 Da.

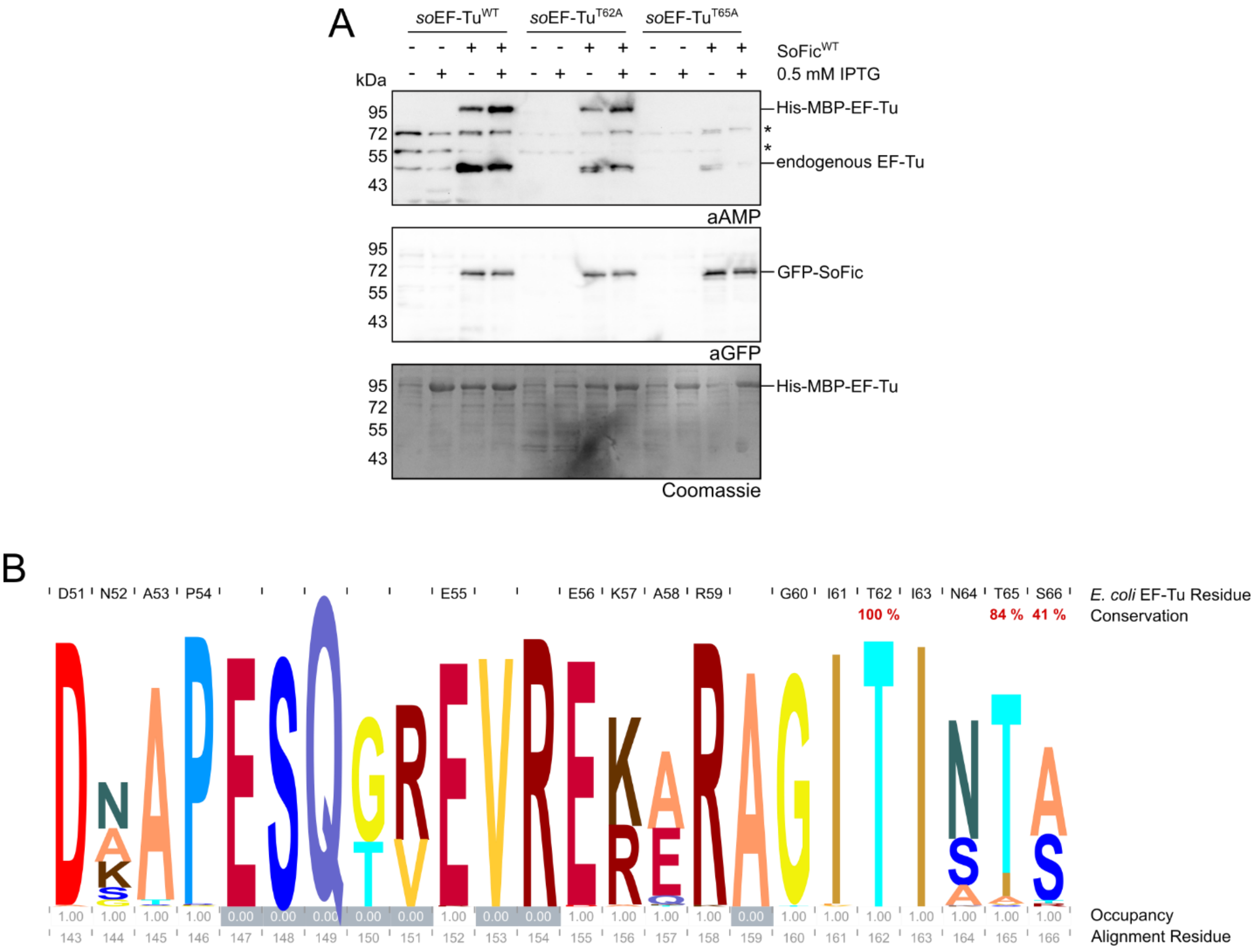

**Figure S6: AMPylation of EF-Tu is conserved across species.**

**A.** Western blot analysis of culture samples after co-expression of GFP-tagged SoFic$^{WT}$ with His$_6$-MBP-tagged *S. oneidensis* EF-Tu and modification site mutants thereof. Culture samples were normalized by OD$_{600nm}$ after cultivation without or with 1 mM IPTG at 22 °C overnight. After detection with an anti-AMP antibody, the membranes were stripped and detected again with an anti-GFP-antibody to control for SoFic expression. Residual protein in the SDS-PAGE gels after Western blotting was stained with Coomassie to control for overexpression of the target proteins. Asterisks (*) indicate endogenous background signals in *E. coli*.

**B.** Logo representation of the conservation of the EF-Tu switch-I region (*E. coli* residues D51-T65). Stack height of the letters correlates with the probability of the respective amino acid at this position. The respective sequence of *E. coli* EF-Tu is given on top of the logo, together with the conservation percentage of the tested modification sites T62, T65 and S66. Below the logo, the occupancy of each residue and the residue number in the multiple sequence alignment (MSA) are indicated. An occupancy of zero indicates rare occurrence of this residue among the aligned sequences. The MSA was performed with Clustal Omega [1] based on the procaryotic EF-Tu superfamily (Interpro CD01884, 831 reviewed protein entries)[2, 3]. The representation was generated with Skylign[4].

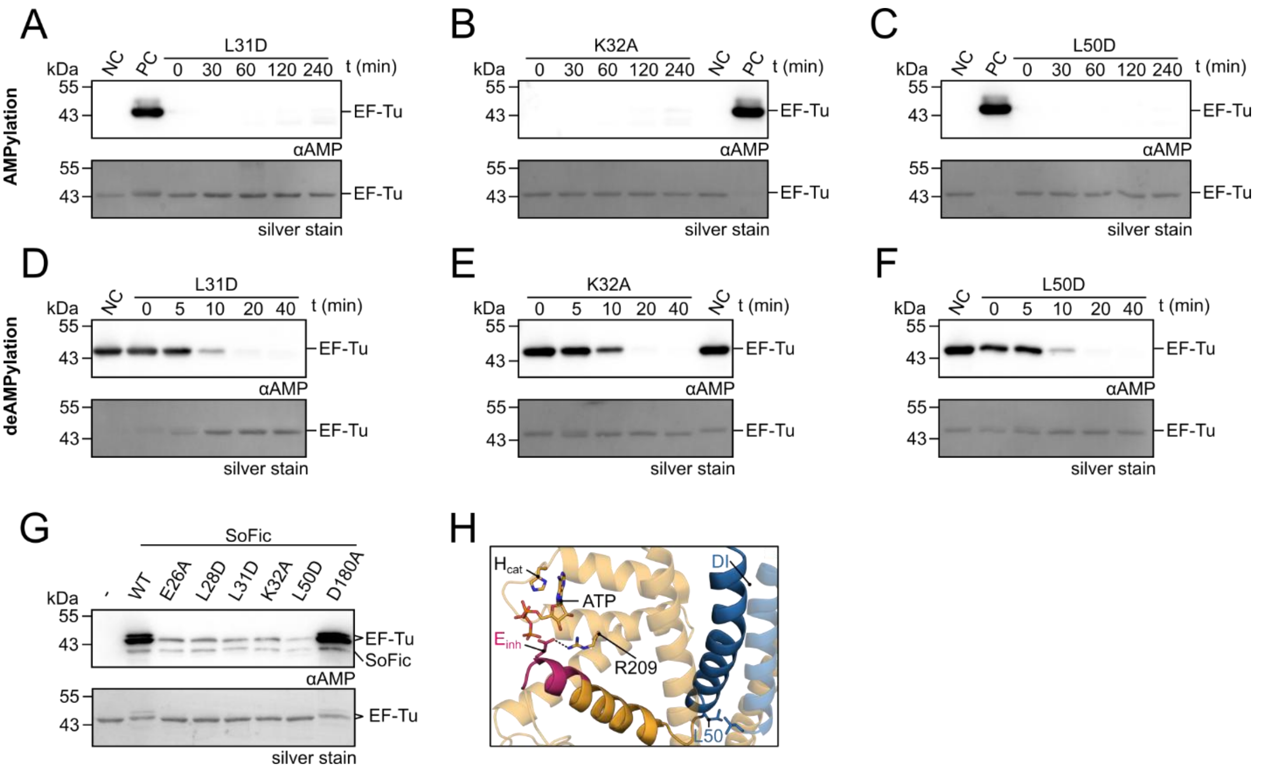

**Figure S7: Dimerization is required for the AMPylation but not the deAMPylation activity of SoFic.**

**A-C.** Western blot analyses of *in vitro* AMPylation reactions with 10 µM *ec*EF-Tu$^{T62A}$ and 0.1 µM of **A** SoFic$^{L31D}$, **B** SoFic$^{K32A}$ and **C** SoFic$^{L50D}$ in the presence of 1.5 mM ATP. At the indicated timepoints, the reactions were stopped by the addition of Laemmli buffer. 250 ng EF-Tu were loaded onto SDS-PAGE for Western blotting, anti-AMP detection and silver staining. NC: negative control without the addition of SoFic (t = 240 min). PC: positive control, 100 % AMPylated *ec*EF-Tu$^{T62A}{}_{AMP}$.

**D-F.** Western blot analyses of *in vitro* deAMPylation reactions with 10 µM *ec*EF-Tu$^{T62A}{}_{AMP}$ and 0.1 µM of **D** SoFic$^{L31D}$, **E** SoFic$^{K32A}$ and **F** SoFic$^{L50D}$. At the indicated timepoints, the reactions were stopped by the addition of Laemmli buffer. 250 ng EF-Tu were loaded onto SDS-PAGE for Western blotting, anti-AMP detection and silver staining. NC: negative control without the addition of SoFic (t = 40 min).

**G.** Western blot analysis of *in vitro* AMPylation reactions with 10 µM *ec*EF-Tu$^{WT}$ and 0.1 µM of the indicated SoFic dimerization interface mutants in the presence of 1.5 mM ATP after incubation at 15 °C overnight. 250 ng EF-Tu were loaded onto SDS-PAGE for Western blotting, anti-AMP detection and silver staining.

**H.** Close-up view of the connection between the dimerization interface (DI, blue) and the inhibitory motif (magenta) of SoFic. Residue L50 at the intersection between both helices, the inhibitory glutamate (E73, $E_{inh}$) and its salt bridge partner R209, the catalytic histidine (H198, $H_{cat}$) and ATP are shown as sticks. The AMPylation deficiency of SoFic$^{L50D}$ and monomeric mutants indicates that allosteric signaling between L50 and $E_{inh}$ may contribute to the relief of autoinhibition in dimeric SoFic. The SoFic:ATP crystal structure from this study (Fig. 1A, Fig. S8, Table S4) was used for the representation.

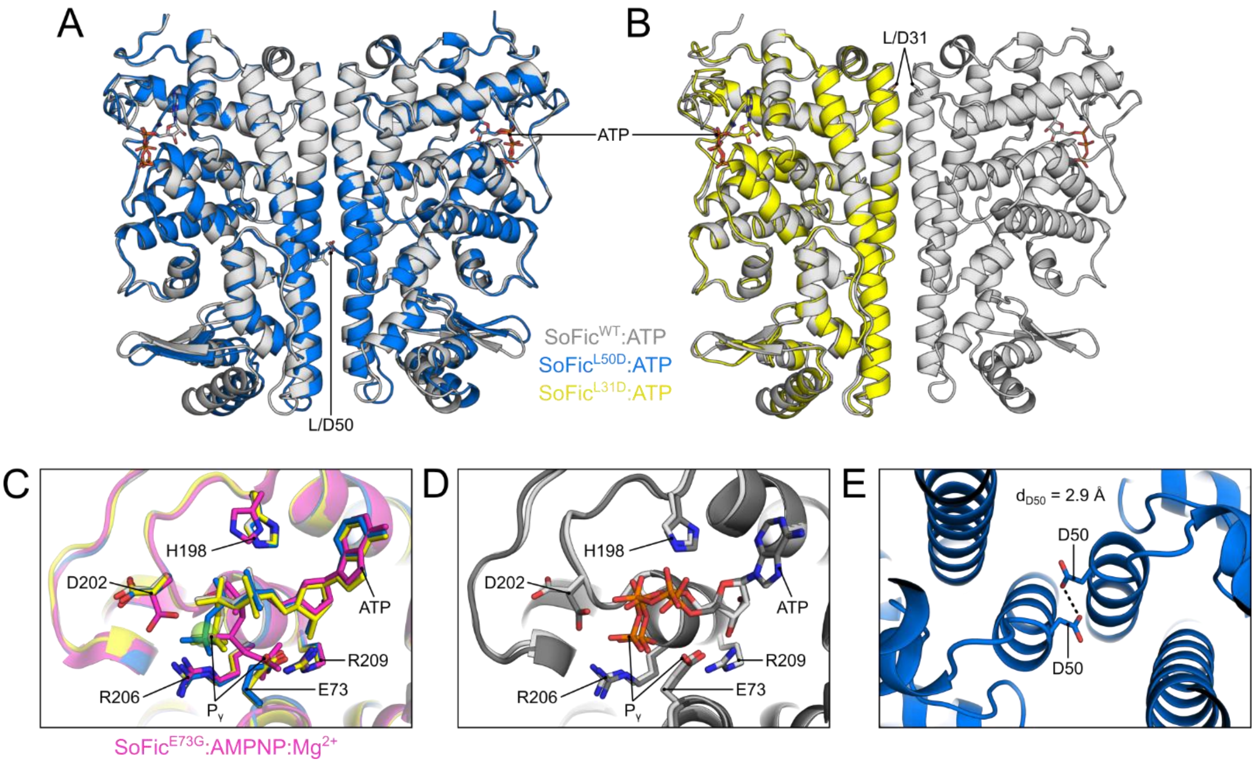

**Figure S8: ATP-bound crystal structures of SoFic$^{WT}$, SoFic$^{L50D}$ and SoFic$^{L31D}$ show minimal differences between monomeric and dimeric mutants.**

**A.** Cartoon representation of SoFic$^{L50D}$:ATP (blue) superimposed to SoFic$^{WT}$:ATP (grey, RMSD = 0.379 Å). ATP is shown as sticks. Chains A and B of both crystals are used for the representation.

**B.** Cartoon representation of SoFic$^{L31D}$:ATP (yellow) superimposed to SoFic$^{WT}$:ATP (grey, RMSD = 0.571 Å). ATP is shown as sticks. Chains A and B of SoFic$^{WT}$:ATP and chain A of SoFic$^{L31D}$:ATP are used for the representation.

**C.** Close-up view of the catalytic center of SoFic:ATP crystal structures. SoFic$^{WT}$, SoFic$^{L50D}$ and SoFic$^{L31D}$ (colored as in A and B) are superimposed to SoFic$^{E73G}$:ATP (pink, PDB 3ZEC[5]). Chain A is shown for all crystals. The triphosphate in the SoFic$^{E73G}$:ATP structure adopts a curved conformation and a $Mg^{2+}$ ion (green sphere) is coordinated by the α- and β-phosphate together with D202. The γ-phosphate of ATP engages forms a salt bridge with R209 of the catalytic motif [5]. In the structures of SoFic$^{L50D}$, SoFic$^{L31D}$, and SoFic$^{WT}$ this γ-phosphate position is occupied by the inhibitory glutamate E73, which forms a salt bridge with R209 instead. Consequently, the γ-phosphate forms a salt bridge with R206 and the triphosphates are arranged in an AMPylation non-competent conformation in all three crystal structures, in line with the absence of electron densities for $Mg^{2+}$ ions.

**D.** Close up view of the catalytic center of the SoFic$^{WT}$:ATP crystal structure. Both dimers in the crystal were superimposed, chains A (light grey) and C (dark grey) are shown. The D202 sidechains of one of two dimers (chains C+D) in the SoFic$^{WT}$ structure point towards the γ-phosphate of ATP, potentially indicating a higher possibility of $Mg^{2+}$ ion coordination for AMPylation catalysis compared to the structures of SoFic$^{L50D}$ and SoFic$^{L31D}$.

**E.** Close-up bottom view of the dimer interface in the SoFic$^{L50D}$:ATP crystal structure. The distance of 2.9 Å between the two D50 sidechains is indicated by a dashed black line. The short distance between two negatively charged residues indicates that the crystallization likely promotes a rigid, stable conformation, that not necessarily reflects the enzyme's conformation in solution.

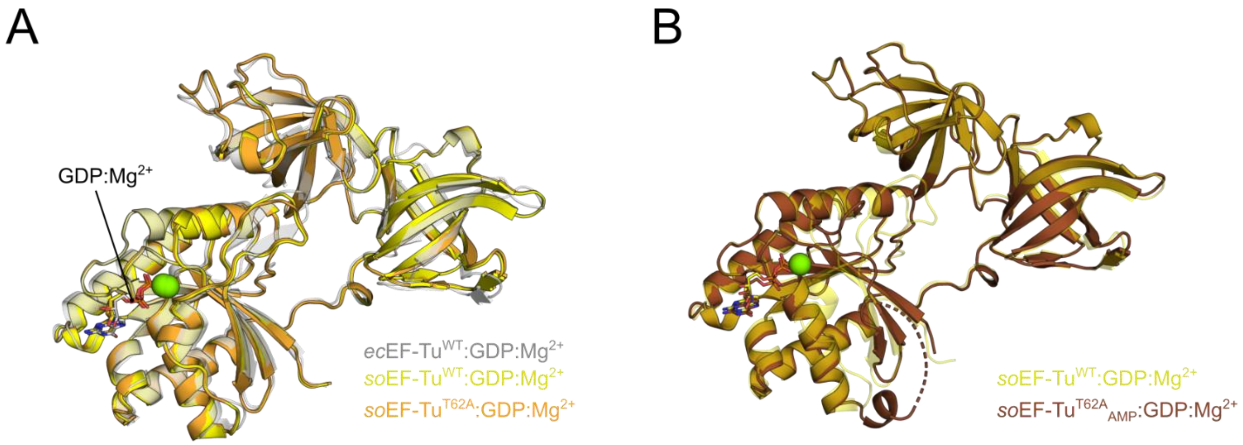

**Figure S9: Crystal structures of unmodified and AMPylated *S. oneidensis* EF-Tu confirm the homology to *E. coli* EF-Tu.**

**A.** Cartoon representation of the unmodified crystal structures of *S. oneidensis* EF-Tu$^{WT}$:GDP:$Mg^{2+}$ (*so*EF-Tu, yellow) and *so*EF-Tu$^{T62A}$:GDP:$Mg^{2+}$ overlaid with *E. coli* EF-Tu:GDP:$Mg^{2+}$ (*ec*EF-Tu, transparent grey, PDB 1EFC[6], RMSD = 0.852 Å and 1.024 Å, respectively). Sticks: GDP, green sphere: $Mg^{2+}$ ion. All three structures adopt the open conformation of EF-Tu, where barrel-I is undocked from the G-domain.

**B.** Cartoon representation of the crystal structure of *so*EF-Tu$^{T62A}$:GDP:$Mg^{2+}$ AMPylated at T65 (brown) overlaid with *so*EF-Tu$^{WT}$:GDP:$Mg^{2+}$ (transparent yellow, RMSD = 0.731 Å). Sticks: GDP, green sphere: $Mg^{2+}$ ion. No electron density was observed for the modified switch-I region of AMPylated *so*EF-Tu$^{T62A}$.

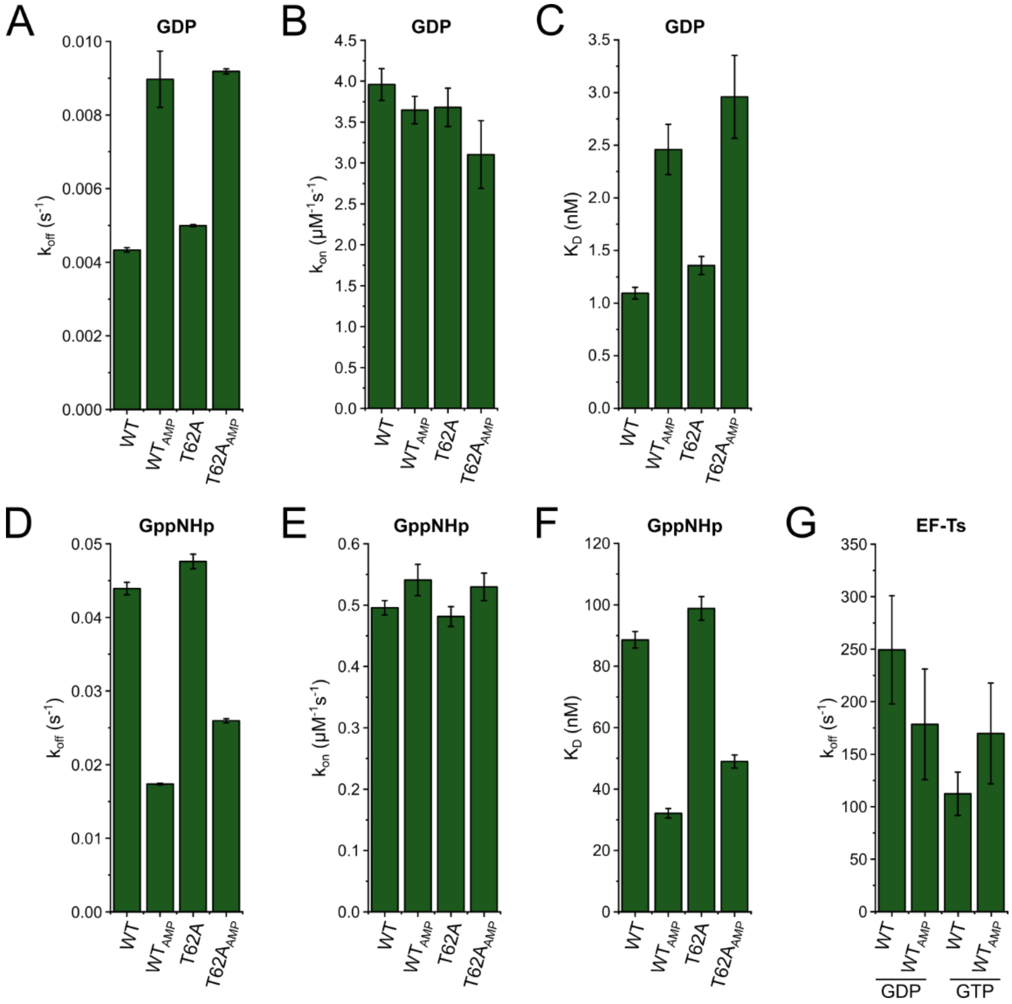

**Figure S10: Stopped-flow kinetics reveal moderate effects of AMPylation on the nucleotide and EF-Ts binding of EF-Tu.**

Stopped flow analyses of *ec*EF-Tu were performed using fluorescent nucleotide derivatives labeled with a 2'/3'-O-(N-Methyl-anthraniloyl)-group (mant-GDP/GppNHp) as reported previously [7]. Förster Resonance Energy Transfer (FRET) from the single tryptophane (W187) of EF-Tu to the mant-group of the labeled nucleotide allows to monitor association and dissociation in a time resolved manner. The dissociation from the nucleotide exchange factor EF-Ts was measured using nucleotide free EF-Tu:EF-Ts complexes. Complex dissociation was induced by the addition of unlabeled GDP or GTP and monitored by the change in Trp fluorescene in EF-Tu [7]. All error bars represent the standard deviation of at least three technical replicates, the data is listed in Table S7.

**A-C.** Dissociation rates ($k_{off}$), association rates ($k_{on}$) and affinity ($K_D$) of unmodified and AMPylated *ec*EF-Tu$^{WT}$ or the T62A mutant for mant-GDP. The increase of $K_D$, i.e. the decreased affinity, is mainly a result of the increased dissociation rates in AMPylated EF-Tu. The $K_D$ values were calculated as the ratio of nucleotide dissociation over association rates.

**D-F.** Dissociation rates ($k_{off}$), association rates ($k_{on}$) and affinity ($K_D$) of unmodified and AMPylated *ec*EF-Tu$^{WT}$ or the T62A mutant for mant-GppNHp. The decrease of $K_D$, i.e. the increased affinity, is mainly a result of the decreased dissociation rates in AMPylated EF-Tu. The $K_D$ values were calculated as the ratio of nucleotide dissociation over association rates. The $K_D$ values of unmodified EF-Tu$^{WT}$ for GDP in **C** and GppNHp in **F** correlate well with the previously published data [7].

**G.** Dissociation rates ($k_{off}$), of EF-Ts from unmodified or AMPylated EF-Tu$^{WT}$ in the presence of unlabeled GDP or GTP. AMPylation may reduce the nucleotide dependent differences in EF-Ts dissociation. However, the relatively large error bars due to a low signal-to-noise ratio in Trp fluorescence measurements complicate a conclusive interpretation of the data.

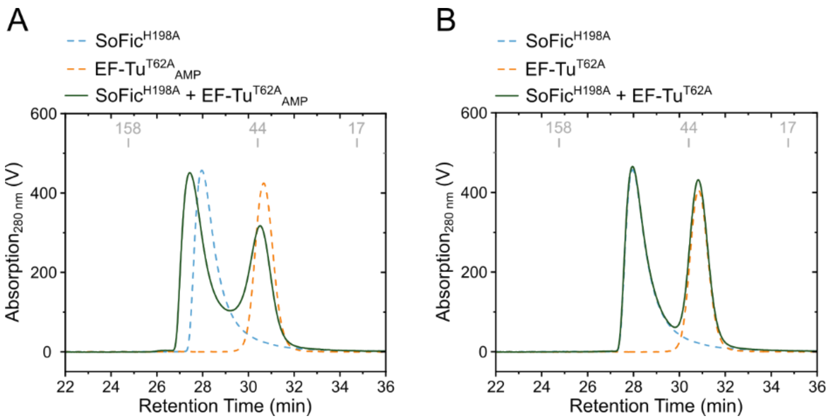

**Figure S11: SoFic$^{H198A}$ and EF-Tu$^{T62A}{}_{AMP}$ co-elute in size exclusion chromatography.**

**A.** Analytical size exclusion chromatography (aSEC) traces of 10 µM SoFic$^{H198A}$ and 10 µM EF-Tu$^{T62A}{}_{AMP}$ alone (dashed blue and orange lines, respectively) and in combination (solid green line). The retention times of reference proteins from a gel filtration standard (BioRad) are indicated in grey.

**B.** aSEC traces of 10 µM SoFic$^{H198A}$ and 10 µM EF-Tu$^{T62A}$ alone (dashed blue and orange lines, respectively) and in combination (solid green line). The retention times of the reference proteins from a gel filtration standard (BioRad) are indicated in grey by their molecular weights in kDa. All traces in **C-F** were baseline corrected.

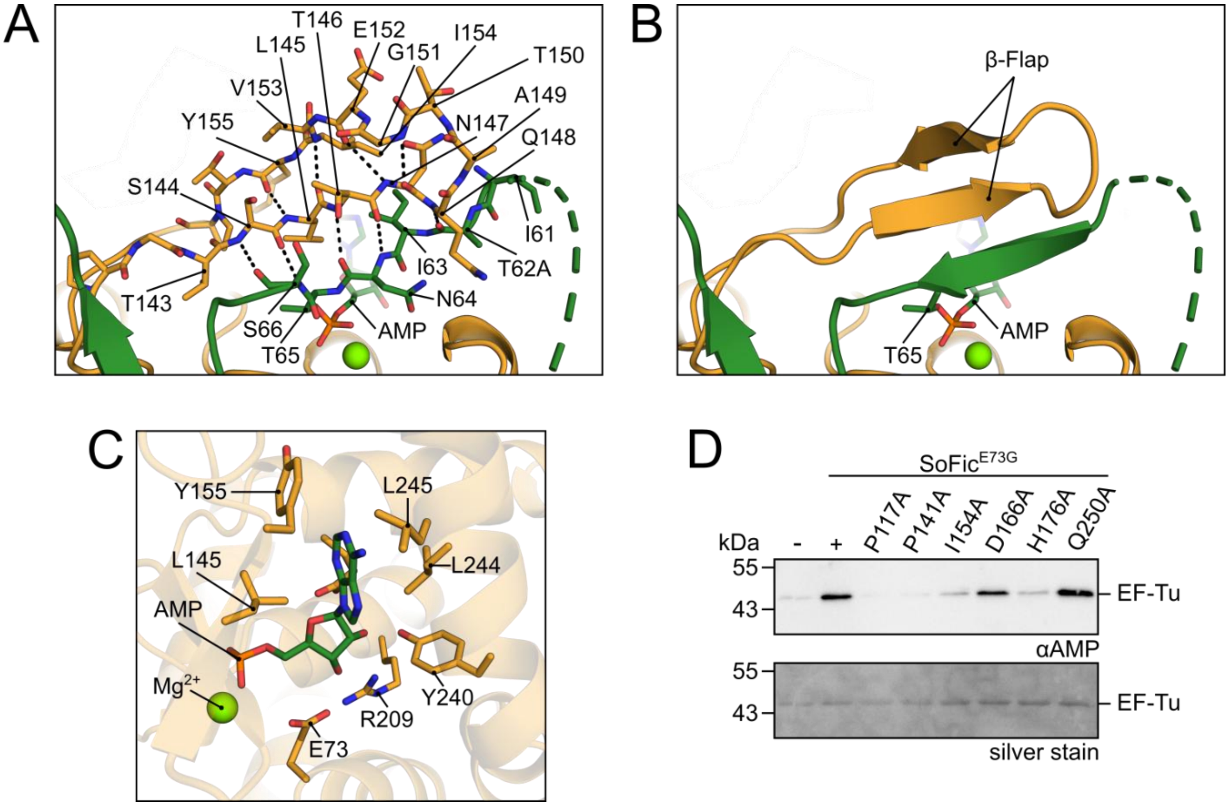

**Figure S12: β-flap interactions and interface validation of the SoFic:EF-Tu complex crystal structure.**

**A.** Stick representation of the AMPylated switch-I region of EF-Tu (green) and the β-flap of SoFic (orange). Backbone interactions are indicated by black dashed lines (distances of 2.8-3.6 Å).

**B.** Cartoon representation of the interaction site in A. The side chain of T65 and the attached AMP moiety are represented as sticks.

**C.** Stick representation of the SoFic residues that form polar contacts with the ribose and hydrophobic interactions with the adenosine of the AMP moiety attached to EF-Tu. The backbone structure of SoFic is shown in transparent cartoon representation. EF-Tu has been omitted for clarity.

**D.** Western blot analysis of *in vitro* AMPylation of 10 µM EF-Tu$^{WT}$ with 0.1 µM His$_{10}$-GFP-tagged SoFic$^{E73G}$ with the indicated additional interface point mutations in the presence of 1.5 mM ATP at 15 °C overnight. 250 ng EF-Tu were loaded onto SDS-PAGE for Western blotting, anti-AMP detection and silver staining. SoFic$^{Q250A}$ was included in the analysis as a control residue which is not involved in the enzyme-target interaction.

### Supplementary Tables

**Table S1: Calculated molecular weights of SoFic mutants in aSEC.** 50 µM of the indicated SoFic mutants were subjected to analytical size exclusion chromatography (aSEC). After alignment to an internal standard, all elution profiles were baseline subtracted and normalized by their maximum intensity. Retention times of the protein peaks were then converted into molecular weights ($MW_{calc}$) using a linear regression of the retention times obtained for a molecular weight standard (BioRad). The standard deviations (SD) represent two replicates using purified SoFic from two independent protein preparations. As a reference, the predicted MW of a SoFic monomer is 42.4 kDa.

| SoFic Mutant | $MW_{calc}$ (kDa) | SD (kDa) |
|---|---|---|
| WT | 89.8 | 0.4 |
| E26A | 53.0 | 0.0 |
| L28D | 46.9 | 0.4 |
| L31D | 47.1 | 0.3 |
| K32A | 47.1 | 0.3 |
| L50D | 79.4 | 1.5 |
| D180A | 85.1 | 0.6 |

**Table S2: DNA Sequences used for binding by SoFic.** The separation of the pDNA into the three fragments F1-F3 is indicated by colors. The DNA sequence of the pDNA corresponds to position 4443980-444279 on the complement strand of the *S. oneidensis* genome (KEGG Genome T00099).

| **Name** | **Sequence (5'-3')** | **Length (bp)** | **GC content (%)** |
|---|---|---|---|
| pDNA | CAAGTAGGCTATCGAGCAGTGTTAAGACTGTTGT CGTCTGCACGTGTATAGATAGTCGTATTATCTATA CACGTGCACTGAATTTGTATAGATATTGGTACTTT CTATACATGTCACTGAACACAAAATGGCAGCCCT AACGCTCCCTTGATGTTGATGCTATCGACATGAG TCTCAAAAACAACCCAAACATGTCGATTAAACTAA GTTTTGTAAACATGTTTTCTTTACATGTCGATTGA ATCGACATAAGCTCACCTCAATCTCTATTTATGTC GATAAAAGAAGGTTTTTTTAAAC | 300 | 36.7 |
| F3 | CAAGTAGGCTATCGAGCAGTGTTAAGACTGTTGT CGTCTGCACGTGTATAGATAGTCGTATTATCTATA CACGTGCACTGAATTTGTATAGATATTGGTA | 100 | 39 |
| F2 | CTTTCTATACATGTCACTGAACACAAAATGGCAG CCCTAACGCTCCCTTGATGTTGATGCTATCGACA TGAGTCTCAAAAACAACCCAAACATGTCGATT | 100 | 42 |
| F1 | AAACTAAGTTTTGTAAACATGTTTTCTTTACATGTC GATTGAATCGACATAAGCTCACCTCAATCTCTATT TATGTCGATAAAAGAAGGTTTTTTTAAAC | 100 | 29 |
| R | ATTCTAAGTGTTGAACGAAATGATTCATGACCTAG AATAAAGTCGTTCTAAAAAATAGATTTTTGTAACTC TCTTCATAAACAATTTGGTGTATCGAAAG | 100 | 29 |

**Table S3: AMPylated peptides identified in LC-MS/MS.** Overview of all AMPylated peptides listed in Supplementary Data Files 2-3. The peptides were identified in LC-MS/MS analyses of SDS-PAGE gel slices corresponding to the endogenous, AMPylated *E. coli* proteins at ca. 45 kDa in Fig. 2A. Indicated are the origin of the gel slice, the identified proteins that contained the AMPylated peptides (indices indicate the amino acid number of the original protein sequence) and the peptide spectra matches (PSMs). The AMPylation sites are given with the ptmRS localization probability in parenthesis [8].

| Sample origin | Identified Protein | AMPylated Peptide | PSMs | AMPylation sites (ptmRS) |
|---|---|---|---|---|
| $SoFic^{WT}$, uninduced culture | EF-Tu | $G_{60}$ITINTSHVEYDTPTR$_{75}$ | 7 | T65 (99) |
| | EF-Tu | $G_{60}$ITINTSHVEYDTPTR$_{75}$ | 3 | T62 (100); T65 (91.9) |
| | 3-oxoacyl-[acyl-carrier-protein] synthase 1 | $L_{54}$DTTGLIDRK$_{63}$ | 1 | K63 (99.8) |
| $SoFic^{E73G}$, induced culture | EF-Tu | $G_{60}$ITINTSHVEYDTPTR$_{75}$ | 7 | T65 (99.4) |
| | EF-Tu | $G_{60}$ITINTSHVEYDTPTR$_{75}$ | 7 | T62(100); T65 (76.7) |
| | 3-oxoacyl-[acyl-carrier-protein] synthase 1 | $L_{54}$DTTGLIDRK$_{63}$ | 1 | K63 (100) |
| | Pyruvate kinase | $G_{157}$VNLPGVSIALPALAEK$_{173}$ | 1 | S164(100) |
| | dITP/XTP pyrophosphatase | $Y_{88}$SGEDATDQK$_{97}$ | 1 | Y88(95.2); S89(95); T94(94.5) |

**Table S4: X-ray data collection and refinement statistics for the crystal structures of SoFic dimerization mutants**. $R_{merge} = \sum_{hkl} \sum_i |I_i(hkl) - \langle I(hkl) \rangle| / \sum_{hkl} \sum_i I_i(hkl)$, where $I_i(hkl)$ is the $i^{th}$ observation of reflection *hkl* and *I(hkl)* is the weighted average intensity for all observations of reflection *hkl*. Statistics for the highest-resolution shell are shown in parentheses.

| | **SoFic$^{WT}$:ATP** | **SoFic$^{L31D}$:ATP** | **SoFic$^{L50D}$:ATP** |
|---|---|---|---|
| PDB Accession | 9TCH | 9TCL | 9TC4 |
| Data Collection | | | |
| Wavelength(Å) | 0.68880 | 0.68880 | 0.68880 |
| Resolution range (Å) | 55.03-2.1 (2.41-2.1) | 46.84-2.66 (2.79-2.66) | 72.92-2.44 (2.48-2.44) |
| Resolution per direction (Å) | 2.09 3.06 2.78 | N.A. | N.A. |
| Space group | P $2_12_12_1$ | P $2_1$ | P $2_12_12_1$ |
| Unit cell dimensions (Å) | 81.92 148.55 161.36<br>90.0 90.0 90.0 | 42.88 58.77 155.17<br>90.0 90.0 90.0 | 81.71 147.97 161.58<br>90.0 90.0 90.0 |
| Total reflections | 727185 (33324) | 156020 (21244) | 843575 (42254) |
| Unique reflections | 53625 (2682) | 22338 (2958) | 73714 (3649) |
| Multiplicity | 13.6 (12.4) | 7.0 (7.2) | 11.4 (11.6) |
| Spherical completeness (%) | 46.3 (6.8) | 99.6 (99.9) | 100 (100) |
| Ellipsoidal Completeness (%) | 86.3 (52.1) | N.A. | N.A. |
| Mean I/σ(I) | 7.9 (2.0) | 6.2 (1.6) | 10.2 (2.4) |
| $R_{merge}$ | 0.32 (1.58) | 0.248 (1.11) | 0.218 (1.23) |
| $CC_{1/2}$ | 0.99 (0.37) | 1.99 (0.78) | 1.00 (0.75) |
| Refinement and model statistics | | | |
| $R_{work}$ | 0.2093 | 0.2400 | 0.1923 |
| $R_{free}$ | 0.2617 | 0.2746 | 0.2300 |
| Number of non-hydrogen Atoms | 12009 | 5872 | 12344 |
| - macromolecules | 11679 | 5763 | 11765 |
| - ligands | 124 | 62 | 124 |
| - solvent | 206 | 47 | 455 |
| RMSD bonds (Å) | 0.003 | 0.002 | 0.002 |
| RMSD angles (°) | 0.529 | 0.484 | 0.520 |
| Ramachandran | | | |
| - favored (%) | 98.43 | 92.47 | 98.97 |
| - allowed (%) | 1.5 | 7.53 | 1.03 |
| - outliers (%) | 0.07 | 0.00 | 0.00 |
| Rotamer outliers (%) | 0.87 | 0.32 | 0.70 |
| Average B-factors ($A^2$) | 40.46 | 47.98 | 48.79 |
| - macromolecules | 40.66 | 48.15 | 49.05 |
| - ligands | 39.82 | 44.53 | 43.02 |
| - solvent | 29.86 | 31.54 | 43.64 |

**Table S5: X-ray data collection and refinement statistics for the crystal structure of AMPylated *E. coli* EF-Tu.** $R_{merge} = \sum_{hkl} \sum_i |I_i(hkl) - \langle I(hkl) \rangle| / \sum_{hkl} \sum_i I_i(hkl)$, where $I_i(hkl)$ is the $i^{th}$ observation of reflection *hkl* and $I(hkl)$ is the weighted average intensity for all observations of reflection *hkl*. Statistics for the highest-resolution shell are shown in parentheses.

| | ***ec*EF-Tu$^{T62A}$$_{AMP}$:GDP:Mg$^{2+}$** |
|---|---|
| PDB Accession | 9TBU |
| Data Collection | |
| Wavelength(Å) | 0.68879 |
| Resolution range (Å) | 60.35 - 1.23 (1.35 - 1.23) |
| Space group | P $2_1$ |
| Unit cell dimensions (Å) | 42.75 76.70 60.38<br>90.0 91.8 90.0 |
| Total reflections | 482670 (20821) |
| Unique reflections | 69905 (3496) |
| Multiplicity | 6.9 (6.0) |
| Spherical completeness (%) | 61.7 (12.2) |
| Ellipsoidal Completeness (%) | 89.0 (41.8) |
| Mean I/σ(I) | 13.7 (1.6) |
| $R_{merge}$ | 0.072 (0.974) |
| $CC_{1/2}$ | 1.00 (0.661) |
| Refinement and model statistics | |
| $R_{work}$ | 0.1735 |
| $R_{free}$ | 0.1908 |
| Number of non-hydrogen Atoms | 3490 |
| - macromolecules | 2963 |
| - ligands | 28 |
| - solvent | 498 |
| - ions | 1 |
| RMSD bonds (Å) | 0.005 |
| RMSD angles (°) | 0.917 |
| Ramachandran | |
| - favored (%) | 2.17 |
| - allowed (%) | 97.83 |
| - outliers (%) | 0.00 |
| Rotamer outliers (%) | 0.31 |
| Avarage B-factors ($A^2$) | 18.60 |
| - macromolecules | 16.68 |
| - ligands | 10.39 |
| - solvent | 30.52 |
| - ions | 11.50 |

**Table S6: X-ray data collection and refinement statistics for the crystal structures of *S. oneidensis* EF-Tu.** $R_{merge} = \sum_{hkl} \sum_i |I_i(hkl) - (I(hkl))| / \sum_{hkl} \sum_i I_i(hkl)$, where $I_i(hkl)$ is the $i^{th}$ observation of reflection *hkl* and *I(hkl)* is the weighted average intensity for all observations of reflection *hkl*. Statistics for the highest-resolution shell are shown in parentheses.

| | ***so*EF-Tu$^{WT}$ :GDP:Mg$^{2+}$** | ***so*EF-Tu$_{T62A}$ :GDP:Mg$^{2+}$** | ***so*EF-Tu$^{T62A}_{AMP}$ :GDP:Mg$^{2+}$** |
|---|---|---|---|
| PDB Accession | 9T8G | 9T9C | 9TCK |
| Data Collection | | | |
| Wavelength(Å) | 0.97626 | 0.97626 | 0.68879 |
| Resolution range (Å) | 60.47-1.61 (1.73-1.61) | 60.48-1.36 (1.49-1.36) | 52.95-1.68 (1.79-1.68) |
| Resolution per direction (Å) | 2.134 1.603 1.608 | 2.06 1.36 1.39 | 1.720 1.842 1.677 |
| Space group | $P\ 2_12_12_1$ | $P\ 2_12_12_1$ | $P2_12_12_1$ |
| Unit cell dimensions (Å) | 63.49 76.74 98.22 90.0 90.0 90.0 | 63.79 76.92 97.89 90.0 90.0 90.0 | 67.22 73.59 76.26 |
| Total reflections | 606925 (28690) | 741780 (37865) | 506929 (25426) |
| Unique reflections | 45515 (2276) | 62952 (3149) | 37214 (1861) |
| Multiplicity | 13.3 (12.6) | 11.8 (12.0) | 13.6 (13.7) |
| Spherical completeness (%) | 71.8 (18.6) | 60.5 (12.9) | 84.5 (24.1) |
| Ellipsoidal Completeness (%) | 94.7 (61.4) | 91.9 (56.0) | 94.9 (57.1) |
| Mean I/σ(I) | 16.9 (1.4) | 13.9 (1.6) | 14.2 (1.4) |
| $R_{merge}$ | 0.077 (1.621) | 0.10 (1.64) | 0.131 (2.013) |
| $CC_{1/2}$ | 1.00 (0.539) | 1.00 (0.73) | 0.999 (0.580) |
| Refinement and model statistics | | | |
| $R_{work}$ | 0.1784 | 0.1888 | 0.1733 |
| $R_{free}$ | 0.2058 | 0.2194 | 0.2060 |
| Number of non-hydrogen Atoms | 3251 | 3489 | 3144 |
| - macromolecules | 2948 | 2983 | 2844 |
| - ligands | 28 | 28 | 28 |
| - solvent | 274 | 477 | 269 |
| - ion | 1 | 1 | 3 |
| RMSD bonds (Å) | 0.011 | 0.004 | 0.015 |
| RMSD angles (°) | 1.158 | 0.769 | 1.348 |
| Ramachandran | | | |
| - favored (%) | 97.92 | 98.18 | 97.23 |
| - allowed (%) | 2.08 | 1.82 | 2.77 |
| - outliers (%) | 0.00 | 0.00 | 0 |
| Rotamer outliers (%) | 0.31 | 0.31 | 0.32 |
| Average B-factors ($A^2$) | 31.62 | 22.92 | |
| - macromolecules | 30.75 | 21.37 | 30.12 |
| - ligands | 22.09 | 14.05 | 26.72 |
| - solvent | 41.97 | 33.13 | 39.82 |
| - ions | 20.62 | 13.70 | 47.74 |

**Table S7: Nucleotide and EF-Ts binding rates of *ec*EF-Tu determined by stopped-flow kinetics.** Values for the association rates ($k_{on}$) were obtained as the slope of a linear fit ($k_{obs} = k_{on} * c + b$) to $k_{obs}$ over concentration ($c$) plots, SD denotes the standard deviation of the linear fit. The dissociation rates ($k_{off}$) were obtained directly from exponential fits to individual dissociation curves ($y = y_0 + A * e^{-k_{off} * x}$), SD denotes the standard deviation of averaging across three to five exchange reactions. The dissociation constant $K_D$ was calculated as the ratio $K_D = k_{off}/k_{on}$, SD was calculated by error propagation using the SD values of $k_{on}$ and $k_{off}$.

| Nucleotide | EF-Tu | $k_{off}$ (*$10^{-3}$ $s^{-1}$) | SD (*$10^{-3}$ $s^{-1}$) | $k_{on}$ ($\mu M^{-1}s^{-1}$) | SD ($\mu M^{-1}s^{-1}$) | $K_D$ (nM) | SD (nM) |
|---|---|---|---|---|---|---|---|
| GDP | WT | 4.335 | 0.058 | 3.959 | 0.195 | 1.095 | 0.056 |
| | $WT_{AMP}$ | 8.970 | 0.763 | 3.647 | 0.169 | 2.459 | 0.238 |
| | T62A | 4.995 | 0.026 | 3.680 | 0.234 | 1.357 | 0.086 |
| | $T62A_{AMP}$ | 9.186 | 0.067 | 3.104 | 0.413 | 2.960 | 0.394 |
| GppNHp | WT | 43.918 | 0.840 | 0.496 | 0.012 | 88.607 | 2.693 |
| | $WT_{AMP}$ | 17.367 | 0.117 | 0.541 | 0.025 | 32.111 | 1.526 |
| | T62A | 47.590 | 0.982 | 0.482 | 0.016 | 98.835 | 3.878 |
| | $T62A_{AMP}$ | 25.950 | 0.284 | 0.530 | 0.022 | 48.992 | 2.139 |
| **Nucleotide** | **EF-Tu:Ts complex** | **$k_{off}$ ($s^{-1}$)** | **SD ($s^{-1}$)** | | | | |
| GDP | WT | 249.413 | 51.507 | | | | |
| | $WT_{AMP}$ | 178.467 | 52.693 | | | | |
| GTP | WT | 112.334 | 20.623 | | | | |
| | $WT_{AMP}$ | 169.762 | 48.033 | | | | |

**Table S8: X-ray data collection and refinement statistics for the SoFic:EF-Tu$_{AMP}$ crystal structure.** $R_{merge} = \sum_{hkl} \sum_i |I_i(hkl) - (I(hkl))| / \sum_{hkl} \sum_i I_i(hkl)$, where $I_i(hkl)$ is the $i^{th}$ observation of reflection *hkl* and *I(hkl)* is the weighted average intensity for all observations of reflection *hkl*. Statistics for the highest-resolution shell are shown in parentheses.

| | **SoFic$^{H1898A}$:*ec*EF-Tu$^{T62A}$$_{AMP}$:GDP:Mg$^{2+}$** |
|---|---|
| PDB Accession | 9TB0 |
| Data Collection | |
| Wavelength(Å) | 0.97625 |
| Resolution range (Å) | 90.35 – 2.76 (3.14 - 2.76) |
| Space group | P 1 |
| Unit cell dimensions (Å) | 66.84 95.94 100.17<br>112.9 100.6 90.4 |
| Total reflections | 105039 (5504) |
| Unique reflections | 29290 (1465) |
| Multiplicity | 3.6 (3.8) |
| Spherical completeness (%) | 80.9 (7.9) |
| Ellipsoidal Completeness (%) | 88.3 (56.8) |
| Mean I/σ(I) | 6.0 (1.6) |
| $R_{merge}$ | 0.06 (0.63) |
| $CC_{1/2}$ | 1.00 (0.66) |
| Refinement and model statistics | |
| $R_{work}$ | 0.194 |
| $R_{free}$ | 0.229 |
| Number of non-hydrogen Atoms | 11657 |
| - macromolecules | 11536 |
| - ligands | 100 |
| - solvent | 19 |
| - ions | 2 |
| RMSD bonds (Å) | 0.001 |
| RMSD angles (°) | 0.477 |
| Ramachandran | |
| - favored (%) | 96.30 |
| - allowed (%) | 3.56 |
| - outliers (%) | 0.14 |
| Rotamer outliers (%) | 0.40 |
| Average B-factors ($A^2$) | 86.05 |
| - macromolecules | 86.05 |
| - ligands (GDP) | 121.49 |
| - ligands (AMP+Mg) | 58.15 |
| - solvent | 53.73 |
